# A unified genetic perturbation language for human cellular programming

**DOI:** 10.1101/2025.11.20.689421

**Authors:** Austin Hartman, Oliver Takacsi-Nagy, Courtney Kernick, Nicole E. Theberath, Johnathan Lu, Lujing Wu, Michelle Mantilla, Siddhesh Mittra, Alison McClellan, Nicole Johnson, Lina Mohamad, Lesly Castillo-Colin, Farzad Hoque, Alexander Eapen, Andy Chen, Laura M. Moser, Trini Rogando, Anabella Hernandez, Katherine Santostefano, Ansuman T. Satpathy, Theodore L. Roth

## Abstract

Evolution simultaneously and combinatorially explores complex genetic changes across perturbation classes, including gene knockouts, knockdowns, overexpression, and the creation of new genes from existing domains. Separate technologies are capable of genetic perturbations at scale in human cells, but these methods are largely mutually incompatible. Here we present CRISPR-All, a unified genetic perturbation language for programming of any major type of genetic perturbation simultaneously, in any combination, at genome scale, in primary human cells. This is enabled by a standardized molecular architecture for each major perturbation class, development of a functional syntax for combining arbitrary numbers of elements across classes, and linkage to unique single cell compatible barcodes. To facilitate use, CRISPR-All converts high level descriptions of desired complex genetic changes into a single DNA sequence that can rewire genomic programs within a cell. Using the CRISPR-All language allowed for head-to-head functional comparisons across perturbation types in a comprehensive analysis of all previously identified genetic enhancements of human CAR-T cells. Combining CRISPR-All programs with single cell RNA sequencing revealed a greater diversity of phenotypic states, including improved functional performance, only accessible through distinct perturbation classes. Finally, CRISPR-All combinatorial genome scale screening of up to four distinct perturbations simultaneously revealed additive functional improvements in human T cells accessible only through iterative multiplexing of modifications across perturbation classes. CRISPR-All enables exploration of a combinatorial genetic perturbation space, which may be impactful for biological and clinical applications.

## MAIN TEXT

The genetic code stores the information of life. Manipulation of that code offers control over cell behavior, and the ability to genetically manipulate the genome of human cells has enabled dramatic advances in basic human biology, genetic therapies, and the generation of foundational datasets. The development of lentiviral^1^ and retroviral vectors^2^ allowed new genetic material to be randomly yet stably and efficiently introduced, enabling the overexpression or therapeutic replacement of natural human genes^3–6^. Viruses also enabled the programmable knockdown of natural genes via the expression of a short interfering RNA (siRNA) from an RNA Pol III promoter^7^. More recently, the discovery and rapid development of CRISPR systems have further allowed programmable and heritable knockout of genes in human cells^8–10^, often similarly requiring viral introduction of DNA encoding an RNA pol III-driven short guide RNA to target the Cas nuclease. Finally, beyond random gene introductions, pairing a CRISPR-targeted nuclease with a DNA template suitable for homology directed or alternative repair pathways^11^ has allowed introduction of new DNA sequences into defined sites in the human genome. Targeted integrations enabled correction of single basepair mutations^12–14^ and, with increasingly larger templates, the insertion of full-length genes. The precision afforded by non-viral integration is therapeutically useful in replacing an inherited deleterious allele^14–16^ or inserting a synthetic gene such as a Chimeric Antigen Receptors^17–19^.

This ever-growing diversity of technical methods has made human gene overexpression, gene knockouts, gene knockdowns, or synthetic gene introductions routine in multiple primary human cell types, such as fibroblasts^20^, T cells^13,14,17^, and hematopoietic stem cells^21,22^. Each of these approaches has been coupled with genome-scale pooled screening applications through combination with next generation sequencing. Tailor-made technologies have each focused on a single perturbation type: whole-genome knockout (e.g. CRISPR-KO screens^23–25^), whole-genome knockdown (e.g. shRNA^26–32^, CRISPRi^33–35^, or Cas13 screens^36–38^), whole genome overexpression (e.g. cDNA/ORF library^39–42^ or CRISPRa screens^34,35^), or large-scale synthetic gene or domain insertions (Pooled knockin^43,44^ or domain^45–51^ screening). A handful of these single-perturbation-type pooled genetic screening approaches have also been linked with complex single cell readouts, such as gene knockouts (Perturb-seq^52,53^, CROP-seq^54^), gene overexpression (OverCITE-seq^42^), or synthetic gene introductions (PoKI-seq^43,44^). Despite these advances, current gene perturbation methods access a limited portion of the possible perturbation space^55^. Namely, current methods are not cross-compatible, are limited to perturbing a few genes at a time, and most often focus on a single class of genetic perturbation^56–59^. Some cross-perturbation-type genetic perturbation technologies have been described, but are not compatible with pooled screening (such as viral gene overexpression with CRISPR base editors or knockouts^60–63^), are largely limited to only testing two gene perturbations at time (e.g. orthogonal CRISPRa + CRISPRi/etc approaches^64–72^, mRNA based CRISPR nuclease + base-editor methods^73,74^, knockin + knockout methods^75,76^), and/or are not compatible with single cell sequencing.

There remains a large gap in our ability to perform generalized genetic perturbations in human cells, whereby multiple complex genetic perturbations, regardless of perturbation type, could be made at genome-scale. Such a generalized genetic perturbation language would aid in genetic therapeutic development, complex mechanistic interrogation of biology, large dataset and cellular model generation^77^, and unification of disparate genetic perturbation technologies^78^. Enhancing the efficacy chimeric antigen receptor (CAR) T cells against human cancers is a challenge that could benefit from a standardized genetic perturbation methodology. While durable remissions have been mediated by CAR T cells in patients with hematologic malignancies, success against solid tumors has been limited^79–81^. Extensive single-perturbation-type studies in primary human T cells have been conducted to identify gene knockouts^82^, gene knockdowns^35^, gene overexpression candidates^83^, synthetic genes^43^, and synthetic binders or signaling domains^47^ to generalize CAR T cell therapies across indications. Yet, scalable, direct comparisons of candidate targets or modifications across different perturbation classes, let alone comprehensive assessment of all proposed genetic enhancements, remains technically infeasible.

Here, we describe the development and application of a new genetic perturbation technology - CRISPR-All - representing a unified genetic perturbation language natively developed in primary human cells to write arbitrarily complex genetic perturbations across type and scale. CRISPR-All allows for any type of genetic perturbation, in any combination, to be examined simultaneously, in high-throughput pooled screens, similar to the perturbation-agnostic genetic exploration undertaken by natural germline evolution or by somatic cells during cancer transformation. We designed a standardized architecture for each major class of perturbation, as well as a standardized syntax for combining these architectures in a single DNA sequence. Crucially, every element is associated with a unique DNA barcode specifying the exact identity of each element included in the construct, enabling large-scale pooled screens, including single cell RNA sequencing. This single DNA sequence, encoding arbitrary combinations of genetic perturbations, can be integrated into a cell’s genome to yield a cell with the desired array of genetic changes. CRISPR-All represents a genetic perturbation language, analogous to computational programming languages, whereby a plain text description of a desired complex array of genetic perturbations can be converted into a high level, standardized syntax that maps to low level genetic code, which when input into a cell’s genome is acted upon to result in the desired genetic program.

### Molecular architecture for universal pooled DNA knockins

Previous methods for scalable pooled screening of DNA coding sequences have been limited by template switching and recombination during library construction and integration, causing a loss of linkage between a coding sequence and its barcode^84–88^. These issues have limited pooled screening of DNA sequences to either short oligo-length libraries, poorly scalable direct construct sequencing (e.g. long read sequencing), or proximal barcoding methods capable of multiplexing only two sequences^44,89^. Further, previous molecular architectures for gene or domain integration screening in human cells have lacked consistent sequence architectures that enable reusability and interoperability among synthesized sequence elements^90^. We thus initially sought to build a molecular architecture for pooled knockin screening that overcame these key issues. Ideally, a pooled knockin method would feature: (1) fully reusable and interoperable DNA sequence elements, (2) arbitrary combination of any number of DNA sequences into a single construct, (3) full user-control over the final expressed coding sequence, and (4) a single unique barcode that can be captured for simple amplicon sequencing readouts and is compatible with single cell RNA sequencing workflows (**Figure 1A**).

**Figure 1.**
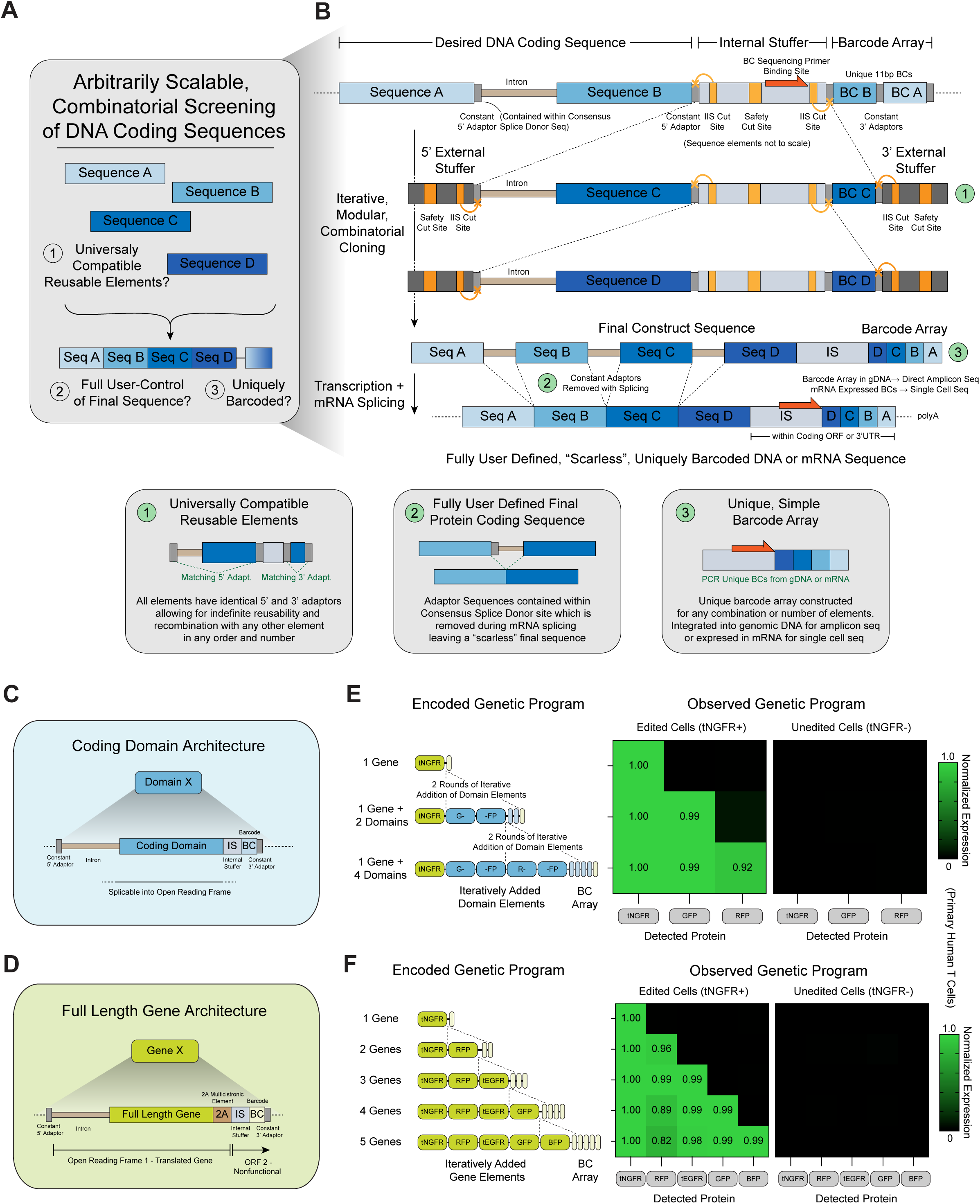
Molecular architecture for universal pooled knockin screening. **(A)** Key unresolved technical constraints for universal pooled gene integration screening include standardized architectures for universally compatible sequence elements, enabling full user control of the final expressed protein sequence(s), and generation of easily detectable unique barcodes for any possible construct combination. **(B)** Key molecular architecture elements to enable arbitrarily complex and combinatorial, pooled knockin sequencing with full user control of the final expressed protein sequence(s). (1) Functional sequences are separated from a unique barcode by a multifunctional “Internal Stuffer” adaptor sequence, with universal adaptors before the functional sequence and internal stuffer (GTAA) and after the internal stuffer and barcode (AGCG). (2) Constant adaptor sequences within the coding frame are encoded within a consensus splice donor sequence of a short intron element, enabling required constant sequences for universal compatibility to be removed from the final coding frame. (3) Iterative cycles of restriction digestions and ligations allow any arbitrary number of DNA elements to be added sequentially to the 5’ end of the Internal Stuffer, with a uniquely identifying barcode array simultaneously being constructed to the 3’ end of the Internal Stuffer. A new Internal Stuffer is inserted within each round, allowing indefinite multiplexing (within ultimate DNA/RNA delivery and integration size constraints). **(C)** Architecture of sub-gene length “Domain” elements for universal pooled knockin screening libraries. For domains integrated at the C terminus of an open reading frame a 2A multicistronic element is added before the Internal Stuffer. **(D)** Architecture of gene length “Domain” elements for universal pooled knockin screening libraries including a 2A multicistronic to allow additional distinct protein products to be sequentially added. **(E)** Flow cytometric validation of successful encoding of up to four Domain elements. Fluorescent reporter genes GFP and RFP were broken into two separate Domain elements each. Fluorescent protein expression measured five days post targeted non-viral integration into the *TRAC* locus of human T cells in n=3 human donors compared to tNGFR only knockin control. **(F)** Validated protein expression of up to five Gene elements using a universal pooled knockin architecture. Fluorescent protein expression measured five days post targeted non-viral integration into the *TRAC* locus of human T cells in n=3 human donors compared to tNGFR only knockin control.

To create universally compatible, arbitrarily combinable elements, we designed an “internal stuffer” / “external stuffer” architecture where each individual sequence element has constant 5’ and 3’ adaptor sequences. When the element is excised from a cloning vector with the external stuffer’s IIS restriction enzyme set, constant sticky end adaptor sequences are created which correspond to the destination sequence’s adaptors created by digestion with the internal stuffer’s IIS restriction enzyme set. These standardized adaptors allow any element to be combined with any other element, while always recreating the internal stuffer so that indefinite numbers of additional elements may be added (**Figures 1B** and **S1A**). While previous iterative combinatorial cloning approaches require constant 4-bp adaptor sequences to be incorporated within the coding frame interspersing each functional sequence^91,92^, we reasoned that these non-desired sequences could be removed from the final coding sequence if they were instead contained within a short intronic region spliced out of the final mature mRNA transcript, leaving only the desired sequence elements behind (**Figure 1B**). Indeed, screening of 24 short introns (52-310 basepairs) identified 11 natural human introns that could be integrated within the coding frame of knocked-in genes in primary human T cells without disrupting expression. In fact, multiple introns could be integrated successfully within one coding sequence (**Figures S1B, 1C**, and **Table S1**). Thus, the constant cloning adaptor sequence could be set to the consensus human “GTAA” splice donor sequence, allowing for a completely scarless coding sequence, even when made up of multiple elements (**Figure 1B**).

To uniquely identify any potential combination of coding sequences, each sequence element to the 5’ end of the internal stuffer is associated with a 11-bp barcode to its 3’ end. Through iterative rounds, a full desired coding sequence is built towards the 5’ end of the internal stuffer while a unique combination of barcodes is built towards the 3’ end (**Figures 1B** and **S1A**). The internal stuffer sequence contains a constant primer binding site for direct PCR amplification of the resulting barcode array from either genomic DNA or mRNA converted into cDNA (**Figure S1D**). Since no scar is present between encoded DNA sequences after splicing, elements could encode for both sub-gene coding domains (**Figures 1C** and **S1E**), or full-length natural or synthetic genes (with a 2A multicistronic element separating them, **Figure 1D**). Indeed, genetic programs constructed using this iterative, intronized, modular approach were successfully integrated into human T cells, with up to five sequence elements, including one gene and four coding domains (tNGFR + GPF1 + GFP2 + RFP1 + RFP2, **Figures 1E** and **S1F**) or five sequential coding genes (tNGFR + RFP + tEGFR + GFP + BFP, **Figures 1G** and **S1G**) showing robust gene expression (>95% in edited cells) by flow cytometry.

### Standardized genetic architecture for protein coding and small RNA element screening

Having demonstrated a potential architecture for universal pooled screening of protein-coding sequences, we next asked whether a similar arbitrarily combinatorial, multiplexable architecture could be applied to non-protein coding sequences such as small RNA effectors (e.g. guide RNAs for gene knockouts and shRNAs for gene knockdowns). Traditional screening approaches using small RNA effectors such as shRNAs or sgRNAs largely rely on expression from an RNA pol III promoter (“U6 Architecture”), with any associated protein components such as selection markers expressed from a separate RNA pol II promoter (**Figure 2A**). Alternatively, both protein coding elements and small RNA elements can be expressed within a single RNA pol II-driven mRNA sequence (“mRNA Architecture”). Small RNA effector elements within the 3’ UTR of an expressed mRNA sequence can be cut out of the mRNA directly such as by a nuclease with gRNA excision capacity such as Cas12a^93^ or by endogenous microRNA processing machinery^94^ (**Figure 2A**).

**Figure 2.**
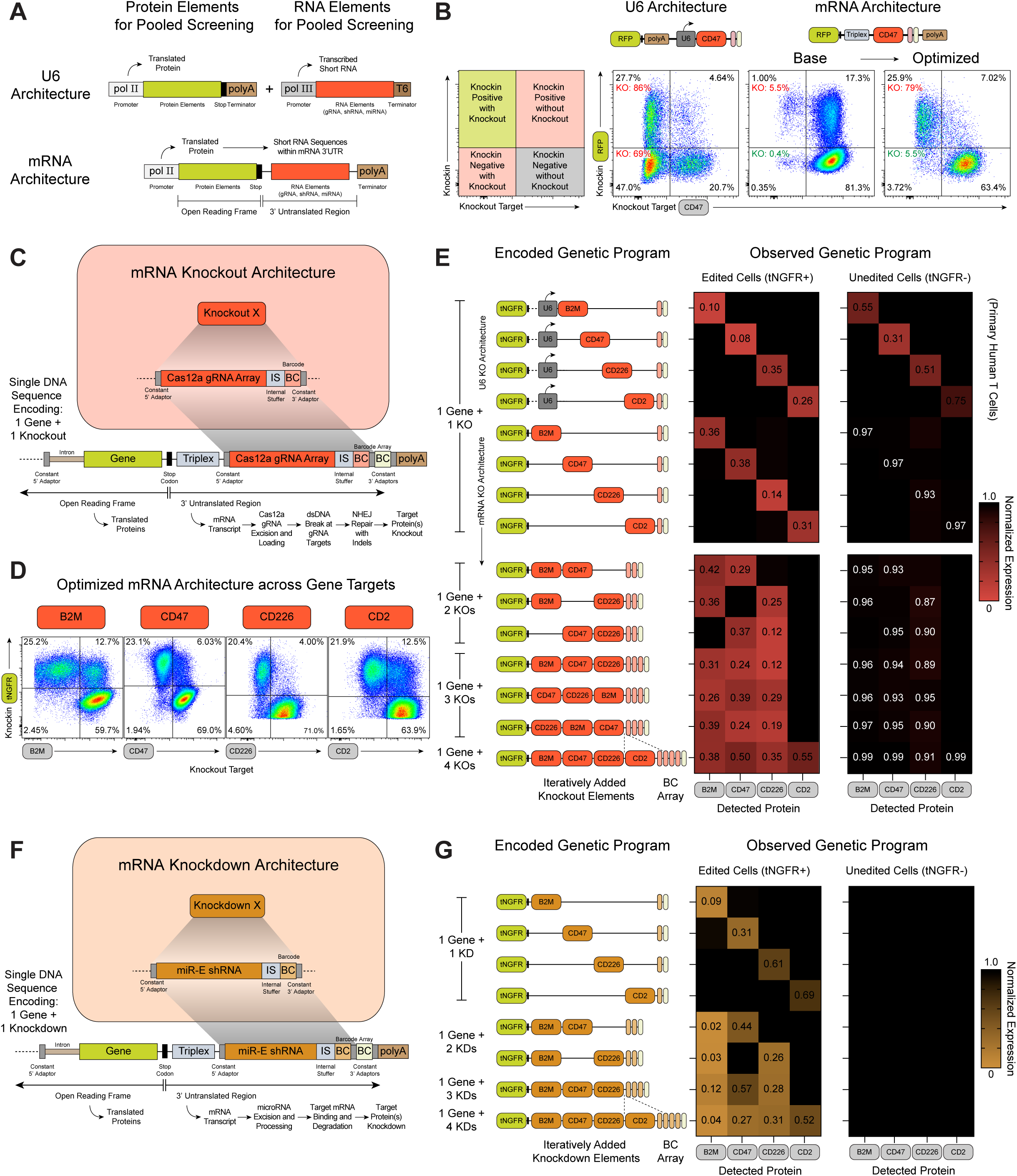
Combinatorial screening of protein and RNA elements. **(A)** Combined protein and RNA element screening (e.g. “Knockin + Knockout” methods) use both Pol II and Pol III promoters in the same construct (“U6 Architecture”). Alternatively, RNA elements can be encoded within non-coding regions of a single RNA Pol II promoter driven expressed mRNA sequence (“mRNA Architecture”). **(B)** Flow cytometric examination of combined Knockin + Knockout performance of U6 vs previously published and optimized mRNA Architectures in primary human T cells by measurement of *CD47* knockout in both knockin positive (tNGFR+) and knockin negative (tNGFR-) cells. Systematic optimization of mRNA Knockin + Knockout architectures (**Figure S2C**) identified an improved mRNA Architecture for cells that resulted in high levels of target protein knockout crucially only in knockin positive cells. **(C)** Final optimized mRNA Architecture for “Knockout” elements integrated with the constant adaptors, Internal Stuffer, and Barcode components from the universal pooled knockin Domain and Gene elements. A Cas12a gRNA array for a target gene is integrated into the 3’UTR of an expressed mRNA transcript (downstream of any protein components). After RNA Pol II driven expression, Cas12a introduced during targeted gene integration (here by electroporation of Cas12a mRNA along with a plasmid DNA HDR Template for targeted non-viral *TRAC* integration into human T cells) excises the gRNAs from the mRNA’s 3’UTR, with the mRNA transcript stabilized by an upstream Triplex RNA stabilization element. Upon binding its target locus, gRNA loaded Cas12a can then induces dsDNA break formation potentially resulting in introduction of frameshift mutations knocking out the targeted protein. **(D)** Efficient and selective Knockin + Knockout of four demonstration target surface receptors using an optimized mRNA architecture expressed in human T cells. **(E)** Flow cytometric measurements of average gene expression of targeted surface receptors in successfully edited (tNGFR+) or uneditined (tNGFR-) human T cells. Across four surface receptor targets, a U6 Architecture showed high levels of knockout in both knockin positive and negative cells, which an optimized mRNA Architecture showed both efficient and selective knockout. mRNA Knockouts could be efficiently multiplexed up to simultaneous knockout of all four targets. **(F)** Optimized mRNA Architecture for “Knockdown” elements, analogous to Knockout elements but with a Cas12a gRNA array replaced by a miR-E shRNA. Endogenous microRNA processing machinery expressed in human cells can excise and process the encoded miR-E shRNA, resulting in small RNA based interference of a target mRNA and resulting protein knockdown. **(G)** Flow cytometric measurements of average gene expression of targeted surface receptors in successfully edited (tNGFR+) or uneditined (tNGFR-) human T cells. Similar to multiplexing of Knockout elements, Knockdown elements showed both efficient and selective knockdown of target surface receptor protein expression, with multiplexing of up to four simultaneous knockdown targets. Knockin and Knockout/Knockdown target protein expression measured by flow cytometry five days post targeted non-viral integration into the *TRAC* locus of human T cells in n=2 human donors (**B, E, E, G**).

We initially examined both U6 and mRNA architectures combined with the internal stuffer-barcode format developed for gene domains and full-length genes. We combined knockin of both an RFP gene and a Cas12a gRNA array targeting the endogenous surface receptor CD47, with the gRNA array expressed either by a Pol III U6 promoter or within a Pol II-expressed mRNA 3’UTR. In human T cells that successfully integrated the U6 architecture (“Knockin Positive”, RFP+), 86% of cells successfully showed knockout of CD47 (**Figure 2B**). However, 69% of cells that did not integrate the U6 architecture (“Knockin Negative”, RFP-) also showed knockout of CD47 (**Figure 2B**), likely due to episomal gRNA expression from the Pol III promoter from non-integrated plasmid HDR templates (**Figure S2A**). Indeed, experiments with libraries of gRNAs introduced by targeted integration with the U6 architecture showed >90% cellular toxicity, again potentially due to episomal expression of multiple unique gRNAs within each cell resulting in intolerable genotoxicity^95,69^ (**Figure S2B**).

In contrast, initial testing of an mRNA architecture based on prior designs^96,97^ showed less than 1% knockout in unedited cells, likely due to restricted expression of the Cas12a gRNA array to only cells where integration of the knockin template occurred (here the *TRAC* locus) (**Figure 2B**). However, the previously described mRNA architectures also showed minimal (~5%) knockout in successfully edited cells (**Figure 2B**). Through iterative testing of Cas12a gRNA array designs, RNA stabilization elements, and editing parameters (**Figure S2C**), we developed an optimized mRNA architecture for Pol II expression of Cas12a gRNAs that resulted in highly efficient (~80%) gene knockouts but crucially restricted gene knockout to knockin positive cells (~5% observed knockout in unedited cells, **Figure 2B**). This selectivity was further confirmed by the rescue of the 90% toxicity seen with U6-expressed gRNA libraries to only ~20% toxicity when the same library was integrated and expressed using an mRNA architecture (**Figure S2D**).

We next combined the optimized mRNA knockout architecture with the same constant internal stuffer–barcode architecture used for coding domains and genes to enable simultaneous expression of protein-coding and gRNA-coding elements (**Figure 2C**). This multiplexable knockout architecture resulted in efficient and selective gene knockout (60-80% knockout with knockin, 2-5% knockout without knockin) across *B2M*, *CD47*, *CD226*, and *CD2* gene targets^98^ (**Figures 2D** and **S2E**). In contrast, U6-encoded architectures across the same four gene targets showed high levels of knockout in both knockin-positive and knockin-negative cells (74-92% knockout with knockin, 25-69% knockout without knockin) (**Figures 2E** and **S2E**). Multiple gene knockouts could be encoded in the same construct, with one gene knockin plus two, three, or even four gene knockout elements still showing highly efficient (69-86% knockout with knockin) and selective (3-7% knockout without knockin) target gene knockout (**Figures 2E** and **S2F**).

Finally, we sought to examine if the remaining major genetic perturbation mode, gene knockdowns, could also be multiplexed with the same optimized mRNA architecture (**Figure 2F**). shRNA sequences were based on miR-E shRNA designs which can be cleaved from longer RNA sequences by endogenously expressed human microRNA processing machinery^94^. Across *B2M*, *CD47*, *CD226*, and *CD2* gene targets, gene knockdown proved to be efficient (20-90% knockdown with knockin) and again selective to only successfully edited cells (<1% knockdown without knockin)^99^ (**Figures 2G** and **S2G**). Gene knockdown elements could also be iteratively added to achieve multiplexed knockdowns, showing successful (20-90% knockdown with knockin) and selective (<1% knockdown without knockin) target knockdown (**Figures 2G** and **S2G**). gRNA based gene knockout could result in lower protein expression of targets than knockdown by shRNA, but across the four tested surface receptor gene targets the performance of shRNAs was more consistent within knockin positive cells (**Figures 2D**, **S2G** and **S2I**). This may reflect gRNAs that cause in-frame indels rather than coding frame disruption. Addition of excisable RNA effector elements to the 3’ UTR of the edited target gene (here the *TRAC* locus) also did not appear to affect the expression of the protein coding gene element (a tNGFR selection marker) (**Figure S2J**).

### Simultaneous genetic screening across perturbation types

Having developed a standardized multiplexable architectures for full length gene, coding domain, gene knockout, and gene knockdown screening, we reasoned these elements may be arbitrarily interoperable with each other. We termed this unified genetic screening system “CRISPR-All”, with each genetic perturbation class serving as a base function type that could be combined in a standardized syntax with any other (**Figures S3A** and **Table S2**). To thoroughly examine the interoperability of Gene, Domain, Knockout, and Knockdown functions, we generated a comprehensive series of six-function CRISPR-All syntaxes, containing every valid combination of four functions (e.g. 4 Genes, 1 Gene + 2 Domains + 1 Knockout, 1 Gene + 2 Knockouts + 1 Knockdown, etc.) along with two constant Gene functions (a tNGFR reporter and a Chimeric Antigen Receptor) (**Figure 3A** and **Table S3-5**). When integrated into the genome of T cells, the predicted genetic programs encoded by this array of CRISPR-All syntaxes were each faithfully observed, and crucially only in successfully edited cells (**Figures 3A** and **S3B**). CRISPR-All encoded genetic programs could reach large complexities. For example, a desired set of ten genetic perturbations was converted into a 10-function CRISPR-All syntax with 4 Gene, 2 Domain, 2 Knockout, and 2 Knockdown functions, which when encoded in a single, uniquely barcoded DNA sequence and knocked into T cells from multiple human donors, demonstrated successful execution of each of the ten functions (>95% expression of each coding gene, 80-95% knockout of *CD2* and *CD226* targets, 85-95% knockdown of *CD47* and *B2M* targets, and equivalent target cell killing to a single CAR gene integration) (**Figures 3C** and **S3B**).

**Figure 3.**
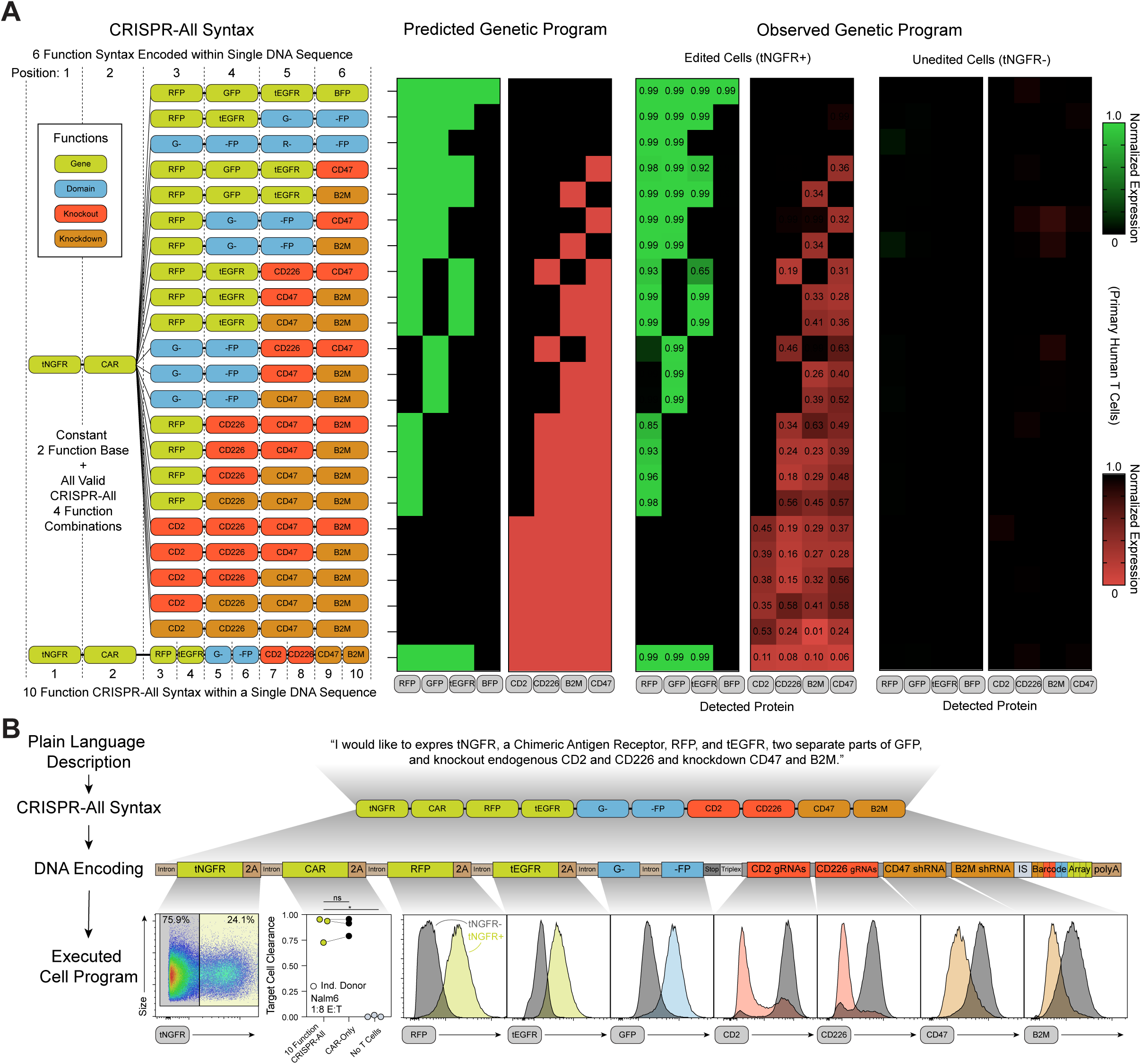
Multiplexed Cross-Perturbation Class Genetic Modification using CRISPR-All. **(A)** The optimized Gene, Domain, Knockout, and Knockdown elements with standardized universal adaptors, Internal Stuffer based multiplexing, and constant barcode architecture for a core set of four Functions for the CRISPR-All genetic perturbation language. All valid four function combinations were generated along with a constant two Gene function base (tNGFR reporter gene and a chimeric antigen receptor). Upon integration of each 6 Function Syntax into human T cells, the observed pattern of protein expression in successfully edited cells (tNGFR+) corresponded to the encoded cross-perturbation class array of genetic programs. The desired genetic program was selectively observed only in successfully edited cells. **(B)** A ten-function program of four Gene, two Domain, two Knockout, and two Knockdown functions described in plain text, along with corresponding CRISPR-All Syntax representation. The resulting full CRISPR-All element encoding converted into a final DNA sequence was integrated into human T cells, resulting in efficient execution of the original described 10 function program. Protein expression measured five days post targeted non-viral integration into the *TRAC* locus of human T cells in n=3 human donors (**A,B**). Functional expression of the CAR (Yescarta) Gene Function demonstrated by a 48 hour *in vitro* killing assay with target Nalm6 cells (1:8 E:T ratio) starting 6 days post editing (**B**). ns = not significant, * = P<0.05, Unpaired t test.

CRISPR-All could be powerful in a context where large amounts of data across major perturbation classes exists, but direct head-to-head comparisons are limited (**Figure 4A**). For example, over the past three decades hundreds of gene knockout targets, gene knockdown targets, gene overexpression targets, CAR signaling domain modifications, full-length CAR architectures, and synthetic gene additions, have been proposed to enhance engineered T cell therapies for cancer^100,101^. We thus hypothesized that CRISPR-All’s ability to screen large scale libraries across perturbation types may enable comprehensive screening and prioritization of genetic targets to improve CAR T cell function.

**Figure 4.**
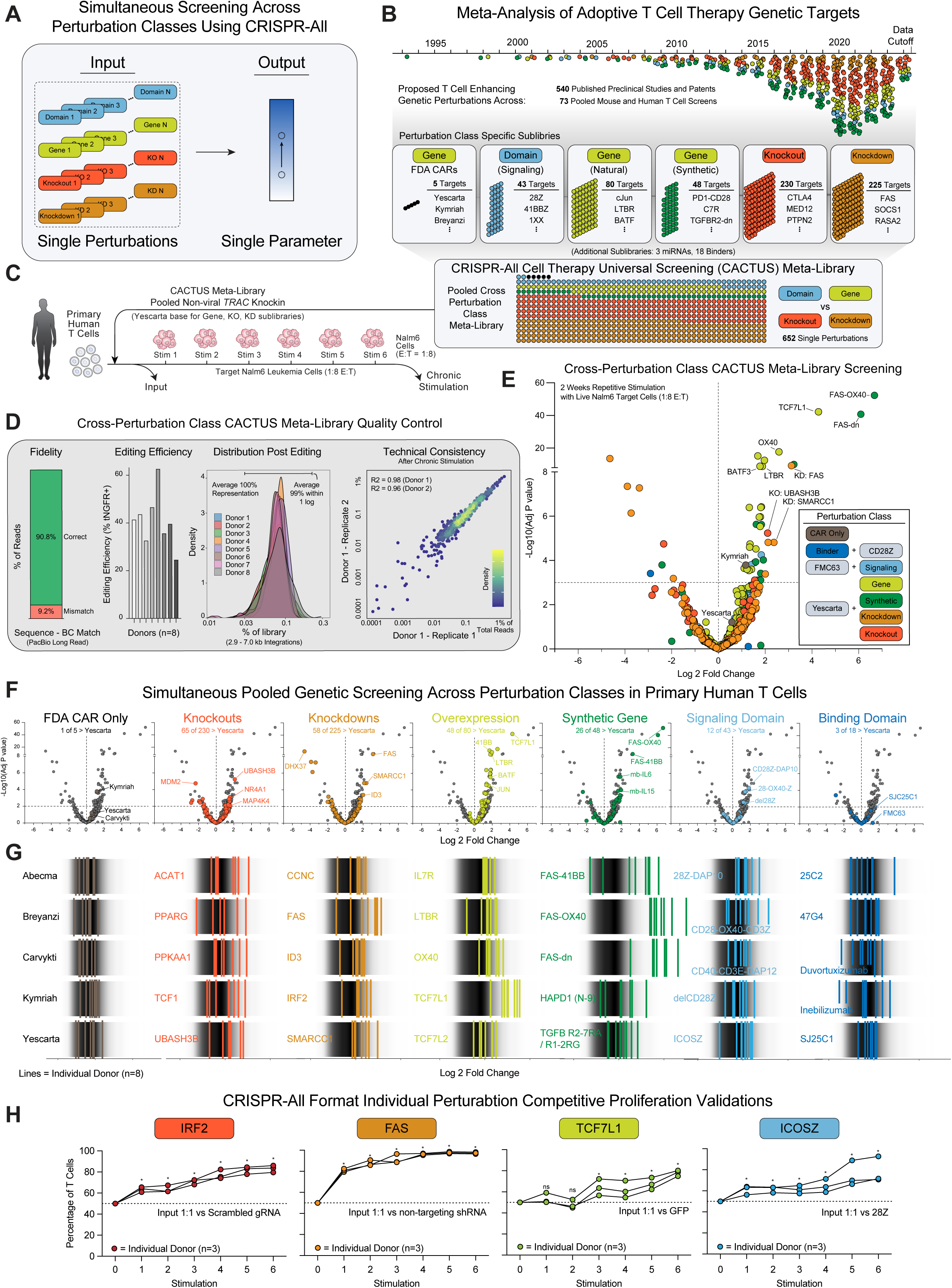
CRISPR-All cellular therapy universal screening meta-library. **(A)** CRISPR-All enables simultaneous pooled screening of diverse perturbation classes head-to-head with scalable barcode based readouts. **(B)** Summary of meta-analysis of adoptive T cell therapy genetic targets. Analysis of 30 years of published literature, patents, and clinical trials with a inclusion cutoff of June 30, 2024 identified 526 publications and 73 pooled screens. Identified targets and top hits from screens were binned into perturbation class specific sub-libraires (CARs, Binders, Signaling Domains, Natural Genes, Synthetic Genes, Knockouts, and Knockdowns) and combined into a single CRISPR-All Cell Therapy Universal Screening (CACTUS) meta-library containing 652 single perturbations across perturbation classes. **(C)** Timeline for CACTUS meta-library repetitive stimulation screen in primary human T cells. CACTUS meta-library integrated T cells were stimulated beginning 6 days post editing with live Nalm6 leukemia cells, reseeding at a 1:8 E:T ratio every 2-3 days for two weeks. Genomic DNA was isolated prior to the first stimulation (“Input”) and two days following stimulation 6 (“Chronic Stimulation”). **(D)** Quantification of CACTUS meta-library fidelity (long read sequencing matching construct’s functional sequence with its barcode within plasmid pool), editing efficiency (n=8 donors), distribution post editing (at Input timepoint, n=8 donors), and technical reproducibility of screen readouts within a single donor (at Chronic Stimulation timepoint, n=2 replicates). **(E)** Results of CACTUS meta-library repetitive stimulation screen in human T cells in n=8 human donors labeled by perturbation class. Functional hits that increased T cell proliferation in the repetitive stimulation assay were identified across perturbation classes. **(F)** CACTUS meta-library repetitive stimulation screen results labeled by perturbation class. **(G)** Donor reproducibility across selected constructs across perturbation classes (each line represents an individual donor’s screen results) **(H)** Validation of increased proliferation in CACTUS meta-library repetitive proliferation screen hits with individual CRISPR-All format perturbations. Across nominated Knockout, Kockdown, Natural Gene overexpression, and CAR Signaling Domain hits, identified targets increased proliferation in individual competitive head-to-head repetitive stimulation validation assay with 1:8 E:T stimulation with Nalm6 target cells (against indicated control constructs). ns = not significant, * = P<0.05, Unpaired t test, n=3 donors.

We first conducted a meta-analysis of adoptive T cell therapy genetic targets nominated in published manuscripts or patent filings identified since 1990 (**Figures 4B** and **S4A**). We identified 613 published preclinical or clinical-stage studies and patents (**Figures S4B, S4C**, and **Table S6**) that nominated specific genetic targets (knockouts, knockdowns, or overexpression) or DNA sequences (Full length CARs, CAR Binders, CAR Signaling domains, or synthetic genes such as chimeric switch receptors) to enhance T cell function. Further, we compiled a second database of over 75 published arrayed or pooled mouse and human T cell screens that sought to identify genetic targets for T cell functional enhancement. These mostly comprised genome-scale CRISPR knockout screens but also included knockdown, overexpression, and synthetic gene and domain screens (**Figure S4B** and **Table S7**). We compiled genetic targets and top screen hits from this meta-analysis into six perturbation-class-specific sub-libraries for FDA approved full length CARs (5), CAR signaling domains (43), natural gene overexpression targets (80), synthetic genes (48), natural gene knockout targets (230), and natural gene knockdown targets (225) (**Figures 4B** and **S4D**). These sub-libraries were combined into a single CRISPR-All format “meta-library” with 636 individual members, that we termed the CRISPR-All Cell Therapy Universal Screening (CACTUS) meta-library (**Figures 4B** and **S4E**; **Tables S8-15**).

To examine CRISPR-All’s ability to effectively screen large libraries of genetic perturbations across perturbation types, we cloned the CACTUS meta-library into a non-viral *TRAC* HDR template vector (**Figures S4F, S4G**, and **Table S16**) and integrated the library into human T cells from eight unique healthy donors (**Figure 4C**). The CRISPR-All formatted CACTUS meta-library demonstrated high individual construct fidelity by long read sequencing (>90% faithful construct-barcode association), efficient integration into human T cells across donors (average ~40% editing efficiency), and tight library distribution (100% representation with 99% of the library within one log) after integration (**Figures 4D** and **S5A-S5D**). We then screened CACTUS meta-library edited T cells in a repetitive stimulation assay, which is commonly used in pre-clinical engineered T cell studies to test proliferation and/or persistence in the setting of T cell exhaustion^44,46,47,102–105^. Here, T cells are repetitively challenged with stress-doses of target live CD19+ BCMA+ Nalm6 leukemia cells every 2-3 days for 6 total rounds of stimulation^106^ (**Figure 4C**). Genomic DNA was isolated before beginning the assay (“Input”) and after two weeks of stimulation (“Chronic stimulation”), and CRISPR-All-formatted barcodes were amplified and sequenced (**Figure S4G**). We gauged the technical consistency of CRISPR-All screening within individual human donors using technical replicates of chronic stimulation screens, which revealed CRISPR-All screening using the CACTUS library was highly reproducible (R2 = 0.96-0.98) (**Figure 4D**).

Given the high fidelity, efficiency, and reproducibility of CRISPR-All-formatted screens, we next asked whether we could simultaneously conduct comprehensive head-to-head functional comparisons of all perturbations identified in our meta-analysis. Indeed, analysis of chronic stimulation screens of the CRISPR-All format CACTUS meta-library in eight human T cell donors revealed enriched gene targets compared to CD19-28ζ (Yescarta) control CAR T cells across each major perturbation class tested, including 65 KOs, 58 KDs, 48 natural genes, 26 synthetic genes, 12 CAR signaling domains, 3 binder domains, and 1 full length FDA approved CAR (Kymriah) (**Figures 4E-4G**, **S5E**, and **Table S17**). CACTUS meta-library screening comprehensively characterized functional enhancements in the screened chronic stimulation environment that improved T cell proliferation and even converged on related targets across different perturbation classes. For example, knockdown of the apoptotic receptor FAS^107^ (L2FC of 3.13, −log10 adj P-Value of 9.1), the overexpression of natural genes such as OX40 (L2FC 2.6, −log10 adj P-Value 17.6) and 41BB^108^ (L2FC 1.7, −log10 adj P-Value 15.1), as well as the introduction of synthetic genes that combined elements of both, such as FAS-OX40^44^ (L2FC 6.7, −log10 adj P-Value 52.2) and FAS-41BB^43^ (L2FC 6.1, −log10 adj P-Value 40.6) switch receptors all were identified as chronic stimulation proliferation hits (**Figure S5F**).

While improved proliferation may be expected from a meta-library of T cell genetic enhancements, many of which were previous proliferation screen hits from single-perturbation-class screening methods (**Tables S6-S7)**, CACTUS screens revealed important context dependence. Knockout/knockdown of the RNA-helicase *DHX37* was a lead hit from a pooled TCR screen for CD8 T cell anti-tumor function^109^, but knockdown of *DHX37* (L2FC −4.6, −log10 adj P-Value 13.6) in the tested CD19 CAR chronic stimulation assay identified it as a highly consistent negative regulator of CAR T cell proliferation (**Figure 4F** and **4G**). Further, CACTUS screens also revealed novel target associations. Overexpression of the genes *TCF7L1* (L2FC 4.3, −log10 adj P-Value 42.1) and *TCF7L2* (L2FC 1.8, −log10 adj P-Value 6.4), previously uncharacterized in T cell contexts (but included within the natural gene overexpression library as orthologs of the transcription factor regulator of T cell development and differentiation TCF7^43^, L2FC 0.97, −log10 adj P-Value 2.7), drove highly reproducible increases in proliferation during *in vitro* chronic stimulation, with over 6-fold greater proliferation than previous top natural gene overexpression hits such as *LTBR*^42^ (L2FC 1.9, −log10 adj P-Value 8.7) and *OX40* (L2FC 2.6, −log10 adj P-Value 17.6) (**Figure 4G**). Finally, representative top constructs from each major CRISPR-All perturbation class identified in the pooled CACTUS chronic proliferation screens were successfully individually validated in head-to-head competitive proliferation assays, with significant (p < 0.05) increases in proliferation seen with CRISPR-All format knockout of *IRF2* and *MED12*, knockdown of *FAS* and *EOMES*, overexpression of *TCF7L1* and *OX40*, and with the CAR signaling domain ICOSZ compared to respective controls (**Figures 4H** and **S5G**). Overall, CACTUS meta-library screening allowed for a comprehensive comparison of TCR and CAR T genetic enhancements across perturbation types in a commonly used pre-clinical repetitive stimulation screening assay. The CACTUS library may be applied in diverse assay environments and screening phenotypes with desired TCR or CAR specificities to prioritize optimal genetic enhancements for specific clinical indications.

### CRISPR-All-seq couples diverse perturbations with single cell sequencing

We next sought to examine whether placing CRISPR-All barcodes at the 3’ end of expressed mRNA proximal to a synthetic polyA sequence allowed for association of cross-perturbation class libraries with scRNA-seq cell barcodes (**Figure 5A**). Each class of CRISPR-All perturbation has the same barcode structure that can be pulled down using standard polyA mRNA capture beads (e.g. 10X 3’ single cell workflows). Then, the cDNA associated with unique cell barcodes can be split to sequence both transcriptomes and CRISPR-All barcodes via a custom PCR step (linking single cell barcodes with CRISPR-All perturbation barcodes) (**Figures 5B** and **S6A**). This method (CRISPR-All-Seq) enables simultaneous cross-perturbation class single cell readouts, analogous to individual perturbation class single cell linkage methods such as Perturb-Seq for gene knockouts^52–54^, OverCITE-Seq for natural gene overexpression^42^, or PoKI-Seq for synthetic genes^43^ (**Figure 5B**).

**Figure 5.**
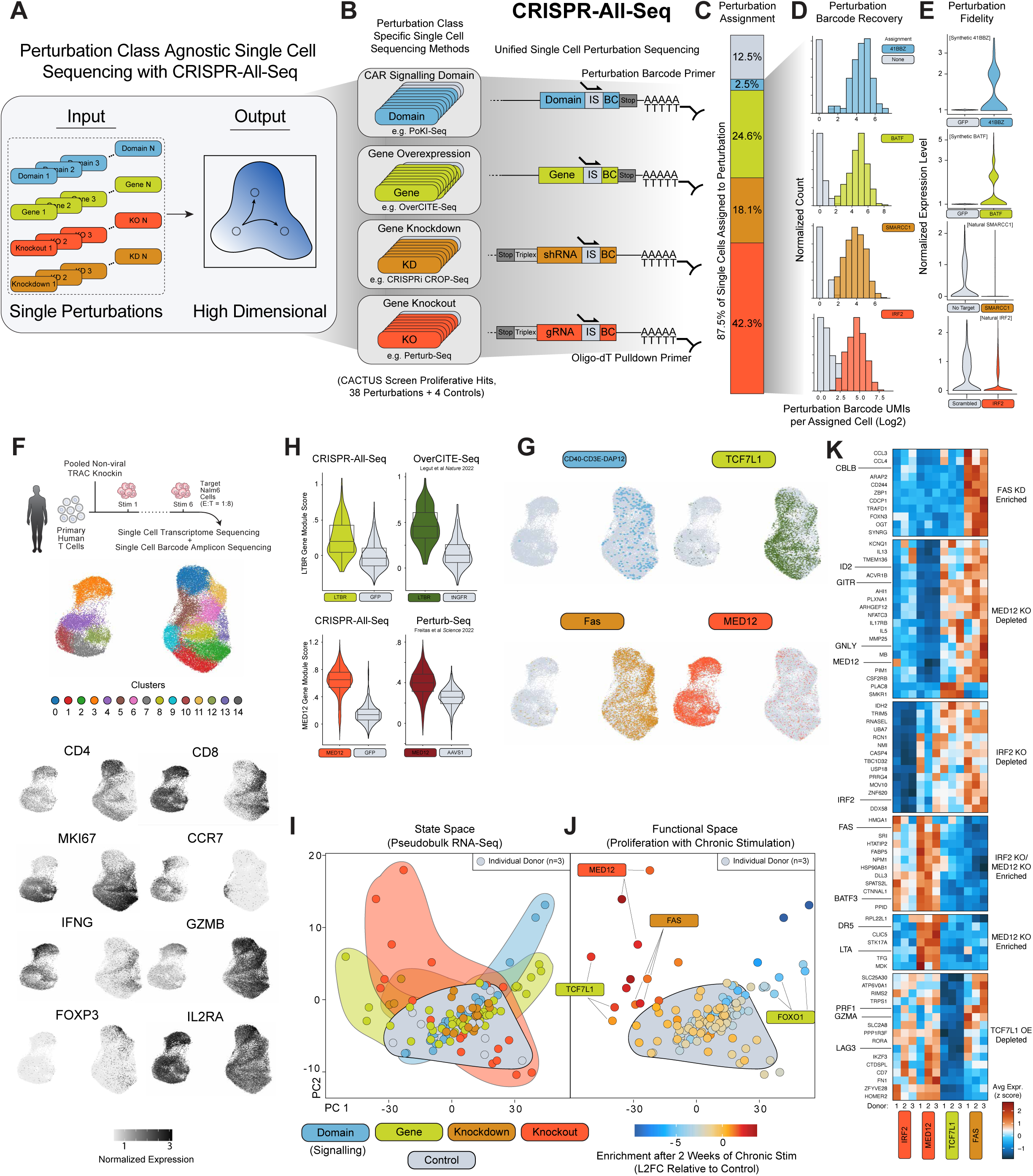
Perturbation agnostic single cell sequencing with CRISPR-All-Seq. **(A,B)** Schematic describing capture and amplification of perturbation barcodes using CRISPR-All-seq. The internal stuffer sequence and perturbation barcodes are contained in the 3’ UTR of the transcript of the knocked-in sequence. The transcripts are captured by conventional poly-dT capture and separately amplified. **(C)** Stacked bar plot of the fraction of cells assigned to each perturbation class with expected skew following a two week repetitive stimulation assay. 12.5% of cells do not have a perturbation assignment due to low barcode UMI counts. 47.1% of cells which are classified as multiplets are subsetted from this display. **(D)** Representative histograms of perturbation barcode UMI counts for a single perturbation of each type on a log2 scale. Cells which are not assigned to a perturbation are shown in grey indicating the background distribution of barcode UMI counts for each displayed perturbation. **(E)** Violin plots displaying perturbation fidelity for cells assigned to each perturbation. The 41BBz and BATF violin plots show expression of codon optimized sequences using kallisto for 41BBz and BATF, respectively, as well as their detection in cells assigned to the GFP control. The SMARCC1 and IRF2 violins show expression of the endogenous SMARCC1 and IRF1 transcripts as quantified by cellranger. Expression values for SMARCC1 and IRF1 are normalized using Seurat with the ‘LogNormalize’ method and 41BBz and BATF values are normalized using the ‘RC’ (relative counts) method. **(F)**Schematic of repetitive stimulation assay where primary human T cells are stimulated 6 times over the course of two weeks with CD19+ Nalm6 cells prior to single cell sequencing. Integrated UMAP embedding of assigned cells colored by unsupervised clustering labels. Additional UMAP embeddings colored by the expression of relevant marker genes. **(G)**UMAP embeddings highlighting four perturbations from each perturbation type. **(H)** Comparison with previously published scRNA-seq datasets describing LTBR overexpression and MED12 knockout showing a shared transcriptomic signature not found in the control cells. Gene module scores are computed using either the CRISPR-All-seq dataset or the previously published dataset (see methods). **(I)** Pseudobulk PCA representation of cells from each donor and perturbation combination. Aggregate counts are batch corrected using limma (see methods). Each donor and perturbation combination is represented by a single point colored by its perturbation type. Background highlights show the embedding space where perturbations from each category are found. **(J)** Embedding is the same as J. Cells are colored by log2 fold-change relative to GFP control abundance. MED12-knockout, FAS knockdown, TCF7L1 overexpression, and FOXO1 overexpression are specifically labelled across 3 donors. **(K)** Heatmap displaying z-scored expression of differentially expressed genes identified for selected perturbations. n=3 donors.

Using a library of 42 genetic perturbations spread across perturbation classes (selected from CACTUS screen hits, **Table S18**), CRISPR-All-seq was able to assign a unique perturbation to 87.5% of identified single cells with a transcriptome that passed QC filters (**Figures 5C** and **S6B**). CRISPR-All-seq was able to clearly identify individual cell’s perturbation barcode, with an average of ~20 perturbation barcode-UMI counts for the assigned perturbation compared to an average of <2 perturbation barcode-UMIs for non-assigned perturbations (**Figures 5D** and **S6C**). Across all four major CRISPR-All perturbation classes, cells assigned to perturbations were orthogonally validated by transcriptomic data. The encoded coding regions for Domain and Gene functions were selectively identified in the single cell transcriptomes of cells assigned the corresponding Domain or Gene CRISPR-All barcode from the separate barcode amplicon sequencing, with a 1.21 log2-fold increase in detection of the expected mRNA sequence relative to the GFP mRNA sequence (**Figures 5E** and **S6D**). Similarly, for Knockdown and Knockout functions, the corresponding transcript was found to be selectively downregulated in assigned cell’s transcriptomes, with an average 0.41 log2-fold decrease in expression detected in Knockdown and 0.35 log2-fold decrease in Knockout assigned cells (**Figures 5E**, **S6E**, and **S6F**). Across cells assigned to each class of perturbation in three separate healthy human donors, single cell transcriptome data showed on average ~8,000 UMIs/cell and ~2,500 unique features per cell (with cell inclusion cutoffs of: unique features > 1000, total UMIs < 40000, and mitochondrial reads < 10%), with a final average of ~1,000 cells assigned to each CRISPR-All perturbation (**Figures S6G-J**).

Analysis of CRSIPR-All-Seq single cell transcriptome data from CD19 CAR-T cells plus one of 42 individual Domain, Gene, Knockdown, or Knockout perturbations after chronic stimulation *in vitro* with Nalm6 target cells showed expression of markers consistent with T cell stimulation and activation (**Figures 5F**). To examine whether CRISPR-All-seq transcriptomic signatures were consistent with existing single cell sequencing methods of individual perturbation classes, we defined gene modules for two CRISPR-All-seq perturbations, LTBR overexpression and MED12 knockout, with previously published single-cell transcriptomic signatures for LTBR (generated by OverCITE-Seq^42^) and MED12 (generated by Perturb-Seq^110^). We found consistent enrichment of LTBR overexpression and MED12 knockout gene modules across single-cell datasets, despite large differences in experimental design and system (**Figures 5H** and **S7A**).

Individual CRISPR-All perturbations showed notable differences in the ability to access specific transcriptomic programs (**Figures 5G**, **S7B** and **S7C**). To examine a high-level view of the transcriptional phenotypes accessed by genetic perturbations across type, we visualized pseudobulked CRISPR-All-seq data from three donors across all 42 individual perturbations (**Figure 5I**). Compared to the phenotypic diversity accessed by control constructs (a 28ζ signaling Domain control, a GFP Gene control, a non-targeting shRNA Knockdown control, and a scrambled gRNA Knockout control, Grey), all four perturbation classes accessed a wider range of transcriptomic phenotypes (**Figure 5I**). Overlaying corresponding functional performance (Log 2 Fold Change in proliferation over the two-week repetitive stimulation assay onto the principle component dimensional reduction revealed that consistently across three human donors, cells with genetic perturbations that accessed the most divergent transcriptomic phenotypes (e.g. MED12 Knockout, FAS Knockdown, TCF7L1 Gene overexpression) corresponded with top functional performance (**Figure 5J**). Perturbations associated with increased proliferation appear to do so, at least in part, by consistently modulating different sets of genes across donors. For instance, we find consistent downregulation of cytotoxic effector genes (*GZMA*, *PRF1, GNLY*) in TCF7L1 OE cells, while the same genes are upregulated in FAS KD cells. We also note significant upregulation of *IFNG* and *SELL* specifically in MED12 KO cells^110,111^ (**Figures 5K** and **S7D**). Overall, CRISPR-All-seq enabled simultaneous cross-perturbation class single-cell sequencing in primary human cells with high fidelity and efficient perturbation assignment regardless of perturbation type.

### Arbitrarily combinatorial pooled genetic manipulations

We next aimed to see whether CRISPR-All could perform multiplexed cross-perturbation class pooled genetic screening (**Figure 6A**). We constructed a four-function combinatorial library of genetic perturbations, with each construct in the library possessing a unique combination of a Domain (CAR signaling domain), Gene (natural or synthetic gene overexpression), Knockout (Cas12a gRNA array), and Knockdown (shRNA), all associated with a uniquely identifying barcode array (**Figures 6B** and **S8A**). Combination of Domain (n=10), Gene (n=16), Knockout (n=8), and Knockdown (n=8) sub-libraries yielded a final library of 10,240 unique four-function combinations (**Table S19**). Analysis of the genome-scale four perturbation function plasmid library prior to targeted integration showed complete representation (100% of the 10,240 x4 function combinations identified) and a tight library distribution (91% of constructs within 1 log), with high sequence fidelity and faithful association of all four barcodes with their unique Domain-Gene-Knockout-Knockdown combination (~60% across library members) given expected levels of template switching during iterative cloning steps (**Figures S8B** and **S8C**).

**Figure 6.**
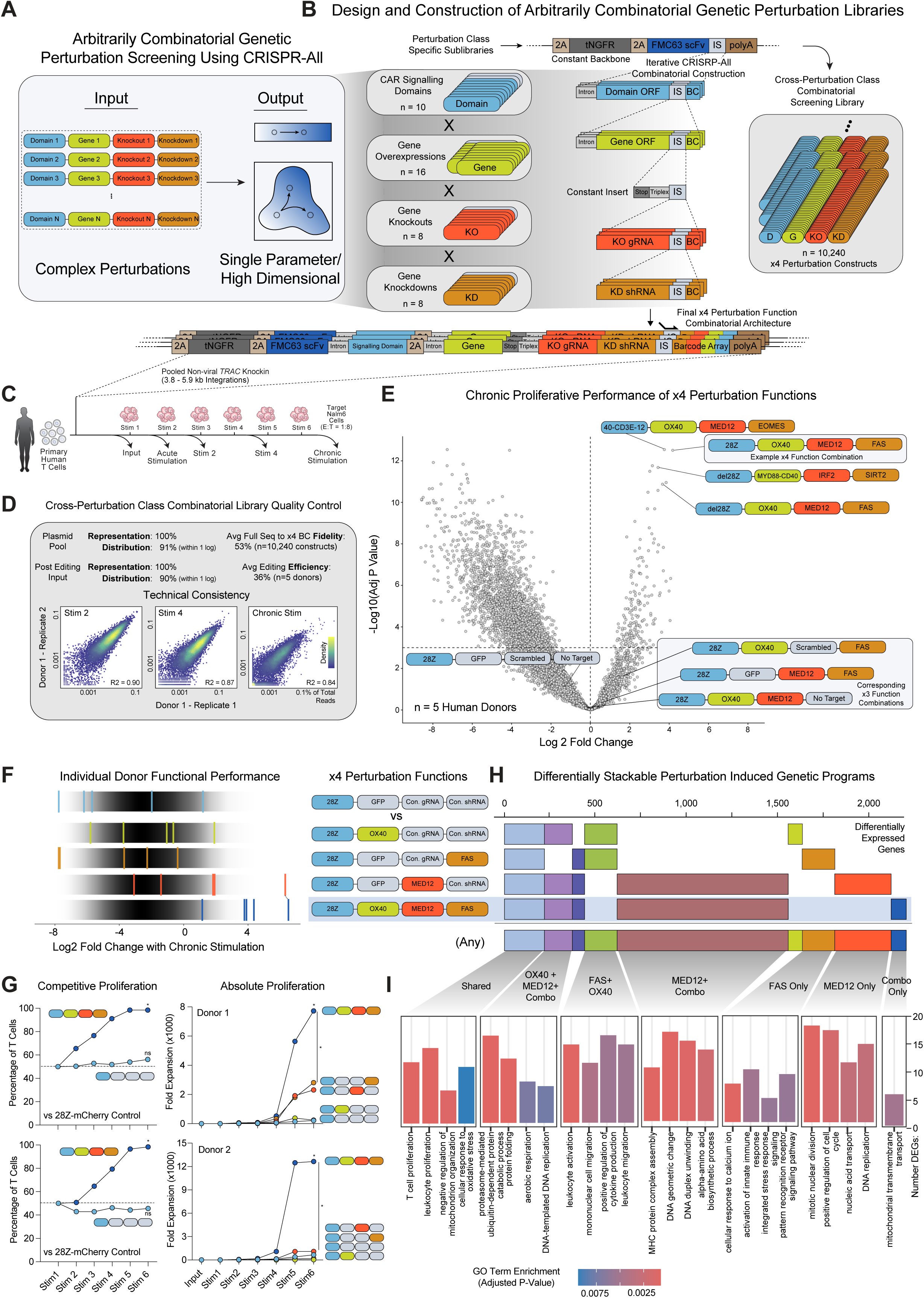
Cross-perturbation class arbitrarily combinatorial genetic screening. **(A)** CRISPR-All enables arbitrary combinations of genetic perturbations to be assayed simultaneously in arrayed and pooled settings with simple single parameter readouts (e.g. proliferation, sorting) or high dimensional readouts (e.g. single cell, functional genomics). **(B)** Iterative cloning workflow to generate a 10,240 member library of four perturbation constructs through combinatorial multiplexing of gene knockdown, knockout, overexpression (natural and synthetic genes), and CAR signaling domain sublibraries. The final construct architecture additionally contains a tNGFR marker gene and an FMC63 CD19-targeting scFv along with four perturbations and a unique barcode array ready for targeted integration. **(C)** Repetitive stimulation screen timeline. Following pooled non-viral knockin to the *TRAC* locus, human T cells with the 10,240 member x4 combinatorial screening library were stimulated with live target Nalm6 leukemia cells at a 1:8 E:T ratio every 2-3 days. **(D)** Quality control metrics of the x4 function combinatorial screening library at the plasmid pool and post-T cell editing phase highlighting representation, distribution, fidelity, editing efficiency, and technical consistency of pooled screening using combinatorial cross-perturbation class CRISPR-All libraries. **(E)** Results of combinatorial x4 perturbation function library abundance screening after chronic stimulation in primary human T cells (n=5 human donors). Top performing constructs were combinations of four perturbation functions, with corresponding constructs possessing x3 or fewer perturbations (containing controls in the other construct positions, such as a scrambled gRNA or a GFP gene) showing decreased abundance. **(F)** Donor consistency of a selected top performing x4 perturbation function combination, 28Z-OX40-MED12-FAS. Across n=5 tested human donors the x4 perturbation function outperformed any individual perturbation on its own in the chronic stimulation abundance screen. **(G)** Individual validation of 28Z-OX40-MED12-FAS x4 perturbation combination in a competitive repetitive stimulation assay (in comparison to a 28Z-mCherry-Scrambled-NoTarget control) showed increased proliferation across n=2 human donors with target live Nalm6 cells (1:8 E:T). A separate 28Z-GFP-Scrambled-NoTarget showed no increase in competitive proliferation. 28Z-OX40-FAS-MED12 x4 perturbation combination also showed increased absolute proliferation in n=2 human donors across repetitive stimulations when individually cultured in comparison to single perturbation constructs. ns = not significant, * = P<0.05, Unpaired t test. **(H)** Bulk RNA sequencing identified patterns of differentially expressed genes between 28Z-OX40-MED12-FAS and individual component perturbations (OX40, MED12, FAS) relative to control edited T cells (28Z-GFP-Scrambled-NoTarget) after two weeks of chronic stimulation. All individual or overlapping sets of >50 DEGs shown in n=3 human donors. **(I)** Gene Ontology term enrichment in selected overlapping sets of differentially expressed genes.

Next, we examined CRISPR-All’s capacity to integrate and reproducibly screen complex perturbation-agnostic combinatorial genetic manipulations. We integrated the genome-scale pooled combinatorial cross-perturbation-class screening library non-virally into the *TRAC* locus of T cells and performed the same chronic stimulation assay as used for the single-perturbation CACTUS meta-library (**Figure 6C**). Across five healthy human donors, the 10,240-member combinatorial perturbation library was efficiently introduced into T cells at high coverage, with an average editing efficiency of 35%, 100% library representation, and 90% of the library within 1 log six days post editing (**Figure 6D**). Side-by-side technical replicates of the chronic stimulation screen in two separate human donors showed high reproducibility (R2 of 0.83-0.90 at Stim 2, 0.31-0.84 at Chronic Stim) (**Figures 6D** and **S8E**). Analysis of construct enrichment revealed consistent top-performers in both acute and chronic stimulation settings, many of which integrated conserved individual elements such as MED12 Knockout, FAS Knockdown, OX40 Gene, and del28ζ signaling Domain (**Figures 6E**, **S8F, S8G**, and **Table S20**). Top performing constructs, such as 28ζ-OX40-MED12-FAS (L2FC 4.0, −log10 adj P-Value 11.9) and CD40/CD3Z/DAP12-OX40-MED12-EOMES (L2FC 3.6, −log10 adj P-Value 12.1) showed high consistency among the five tested human donors (**Figures 6E** and **S8F**). Each perturbation class, in particular the Gene, Knockout, and Knockdown components, appeared to be important for the overall degree of proliferative enhancement seen in top x4 perturbation combinations. Given the inclusion of a control element for each perturbation type during cloning, we could compare top x4 perturbation combinations to constructs where x3, x2 or x1 perturbations were expressed with control elements. Top-performing x4 perturbation combinations were reliant on all four perturbations for maximum enhancement as corresponding x3, x2 or x1 perturbation constructs were enriched less on average or were more variable across human T cell donors (**Figure 6F** and **S9A**).

We individually validated the top-performing four-function perturbation combinations. The 28ζ-OX40-MED12-FAS x4 perturbation combination showed robust individual validation in competitive and absolute proliferation assays, with an average of over 8X greater proliferation over six total stimulations compared to any tested control or single perturbation constructs (e.g. OX40, MED12, or FAS on their own with 28ζ) (**Figure 6G**). A second x4 function perturbation combination, with a CD40-CD3Z-DAP12 Signaling, OX40 Gene, MED12 Knockout, and EOMES Knockdown, similarly showed high donor consistency and functional validation across head-to-head and absolute proliferative assays, with an average 6X greater proliferation than any individual perturbation (**Figure S9B**). Further, both tested x4 function combinations showed improved killing capacity *in vitro* compared to controls, and enhanced chronic killing capacity compared to each individual component perturbation (**Figure S9C**).

Finally, we examined how each individual perturbation component contributed to the overall cell phenotype produced by the x4 combinatorial perturbation construct. Using CRISPR-All’s ability to encode arbitrary single or combinatorial perturbations across perturbation type with a consistent molecular architecture, we performed bulk RNA sequencing for both the 28ζ-OX40-MED12-FAS and CD40/CD3Z/DAP12-OX40-MED12-EOMES x4 perturbation combinations in comparison with each perturbation component individually (**Figures 6H** and **S9D**). The transcriptomic program induced by the combinatorial 28ζ-OX40-MED12-FAS x4 perturbation construct (measured as differentially expressed genes compared to 28ζ signaling domain only) indeed included a base T cell proliferation-associated gene set that was shared across all three individual perturbation components (i.e. OX40 overexpression, MED12 knockout, and FAS knockdown), a metabolic signature from the combination of OX40 and MED12 individual programs, and a large DNA replication-focused program found only in MED12 and the combinatorial x4 perturbation construct (**Figures 6I**). Interference between individual perturbation programs was also observed, such as loss of a migration-focused program found within both FAS and OX40 or a nuclear division program associated with MED12 but not present in the combinatorial 28ζ-OX40-MED12-FAS x4 perturbation construct (**Figures 6I**). Bulk RNA and GO term enrichment analysis for a second validated x4 perturbation construct (CD40/CD3Z/DAP12-OX40-MED12-EOMES) again showed distinct sets of combinatorial gene expression stacking and interference (**Figures S9D** and **S9E**).

## DISCUSSION

CRISPR-All represents a consolidation of diverse perturbation class specific methodologies to produce single or high throughput pooled genetic manipulations, such as CRISPR-KO^23–25^, CRISPRa^33–35^, CRISPRi^33–35^, shRNA^26–28^, cDNA/ORF libraries^39–42^, pooled knockin^43,44^, Cas13^36–38^, or pooled domain screens^45–49^. Many of these methods are multiplexable within their perturbation class, such as dual CRISPR-KO screens^112–114^, multiplexed CRISPRa or CRISPRi, Cas13 screens^37,38^, or two-construct combination pooled knockin screening^44^. CRISPR-All enables genome-scale pooled genetic screening of indefinitely multiplexable, combinatorial genetic perturbations arbitrarily across major perturbation classes. Natural gene overexpressions, synthetic gene introductions, coding or non-coding domains, gene knockdowns, and knockouts can be screened simultaneously against each other in the same pool of cells at the same time, individually or in arbitrary combinations.

We demonstrate CRISPR-All’s unique pan-perturbation type screening capacity in human T cells by generating a “meta-library” comprised of over 30 years of proposed genetic enhancements to drive improved function in TCR and CAR T cell therapies. Combining results from over 600 preclinical studies and over 75 large-scale mouse and human single-perturbation class screens, we generated a CRISPR-All Cell Therapy Universal Screening (CACTUS) meta-library including over 600 distinct genetic enhancements identified by the cellular therapy field, including full-length CAR constructs, CAR binder domains, CAR signaling domains, natural gene overexpressions, synthetic gene introductions, natural gene knockdowns, and natural gene knockouts. Screening of the CACTUS meta-library in a widely used repetitive stimulation pre-clinical development model^44,46,47,102–105^ enabled head-to-head comparison of all prior perturbations in a unified setting. This revealed the most efficient genetic manipulations to drive continued proliferation in exhaustion-inducing *in vitro* chronic stimulation settings, including cross-perturbation class hits manipulating the same biological pathways, context-dependent positive and negative regulators of proliferation, and more powerful functional orthologs of previous hits. Further, combinatorial CRISPR-All screening of up to four perturbation types simultaneously identified multiplexed cross-perturbation class constructs that drove even greater enhancements in proliferation than any previously identified individual perturbation included within the CACTUS meta-library. While we performed these screens in an *in vitro* model of T cell exhaustion with most perturbations coupled to a clinically used CD19-28ζ CAR (Yescarta), future applications of the CACTUS meta-library into more complex *in vitro* and *in vivo* model systems with more diverse functional readouts will reveal ideal tumor indication- and CAR construct-specific functional enhancements to prioritize for limited available slots in clinical trials.

CRISPR-All-seq, the single cell implementation of CRISPR-All, enables high-dimensional phenotypes to be associated with complex perturbation-agnostic genotypes. Indeed, application of CRISPR-All-seq in T cells revealed that the full diversity of potential transcriptomic phenotypes in the examined chronic stimulation assay could only be accessed when screening simultaneously across perturbation types, and that the most diverse transcriptional phenotypes were associated with the greatest functional changes in proliferation. As artificial intelligence-guided cellular modeling and perturbation prediction efforts continue to expand^115–117^, current individual gene knockout or individual gene overexpression single cell data generation workflows may impose limitations. For generation of training data for cellular models, input datasets that are only able to identify transcriptomic states accessible with a single gene’s expression being turned up or down, and which are generated using disparate technical systems susceptible to batch effects, impose fundamental constraints on the ultimate predictive power of cellular models. CRISPR-All’s native capacity to link arbitrarily complex genetic perturbations with single cell sequencing will enable both scalable complex training data generation across a greater range of human cellular states, and rapid predictive testing of proposed complex genetic manipulations regardless of type or number. Indeed, the ability to make arbitrarily complex genetic perturbations at scale may open access to new, non-evolved human cell states that could correlate with improved biologic or therapeutic functions.

More broadly, CRISPR-All in many ways resembles a computational programming language (**Figure 7A, 7B**). CRISPR-All allows a user with limited molecular biology experience to input a desired set of genetic perturbations regardless of perturbation type or number and be returned an “executable” DNA sequence that will result in the desired set of perturbations when delivered (through targeted knockins or other delivery modes) into a cell type of interest.

**Figure 7.**
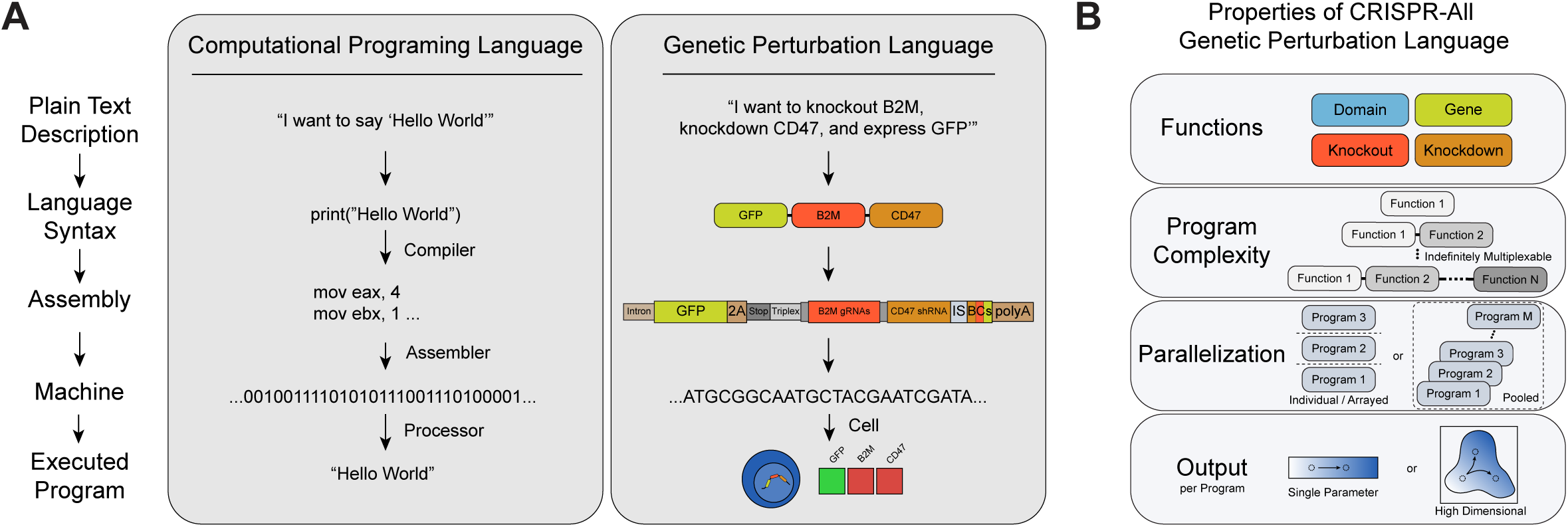
CRISPR-All, a unified genetic perturbation language. **(A)** CRISPR-All serves as a genetic perturbation language, analogous to a computational programming language. CRISPR-All converts a plain text description of a desired set of genetic perturbations into a standard Syntax that maps to successfully lower levels of element encoding and ultimately a raw DNA sequence that can be executed by a cell, abstracting underlying molecular complexity away from the user similar to computational programming languages conversion of high level programming functions through a standardized language syntax into lower level assembly code and ultimately machine readable binary code executable by a CPU. **(B)** Core properties of the CRISPR-All genetic perturbation language. A base set of four perturbation Functions (Gene, Domain, Knockout, Knockdown) can be assessed individually or in arbitrarily multiplexable combinations. CRISPR-All supports parallelization (pooled screening) of any combination of single or multiplexed perturbation Functions simultaneously, with simple amplicon sequencing of genomically integrated barcode arrays for high throughput single parameter readouts. CRISPR-All further natively supports high dimensional single cell readouts linked with any combination of perturbation Functions as barcode arrays are expressed in the 3’UTR of mRNA transcripts.

CRISPR-All unifies disparate genetic discovery technologies to enable defined genetic changes to be made arbitrarily, combinatorially, and scalably, mirroring natural evolution’s exploratory potential to discover new cellular functions and states via diverse genetic perturbations. CRISPR-All will drive fundamental biologic discovery applications across human and model organism cell types, serve as an ideal molecular interface for training data generation and functional prediction testing for large-scale computational cell models, and accelerate development of both integrated and transiently expressed next-generation genetically encoded therapies.

## SUPPLEMENTARY FIGURE LEGENDS

**Supplementary Figure 1.**
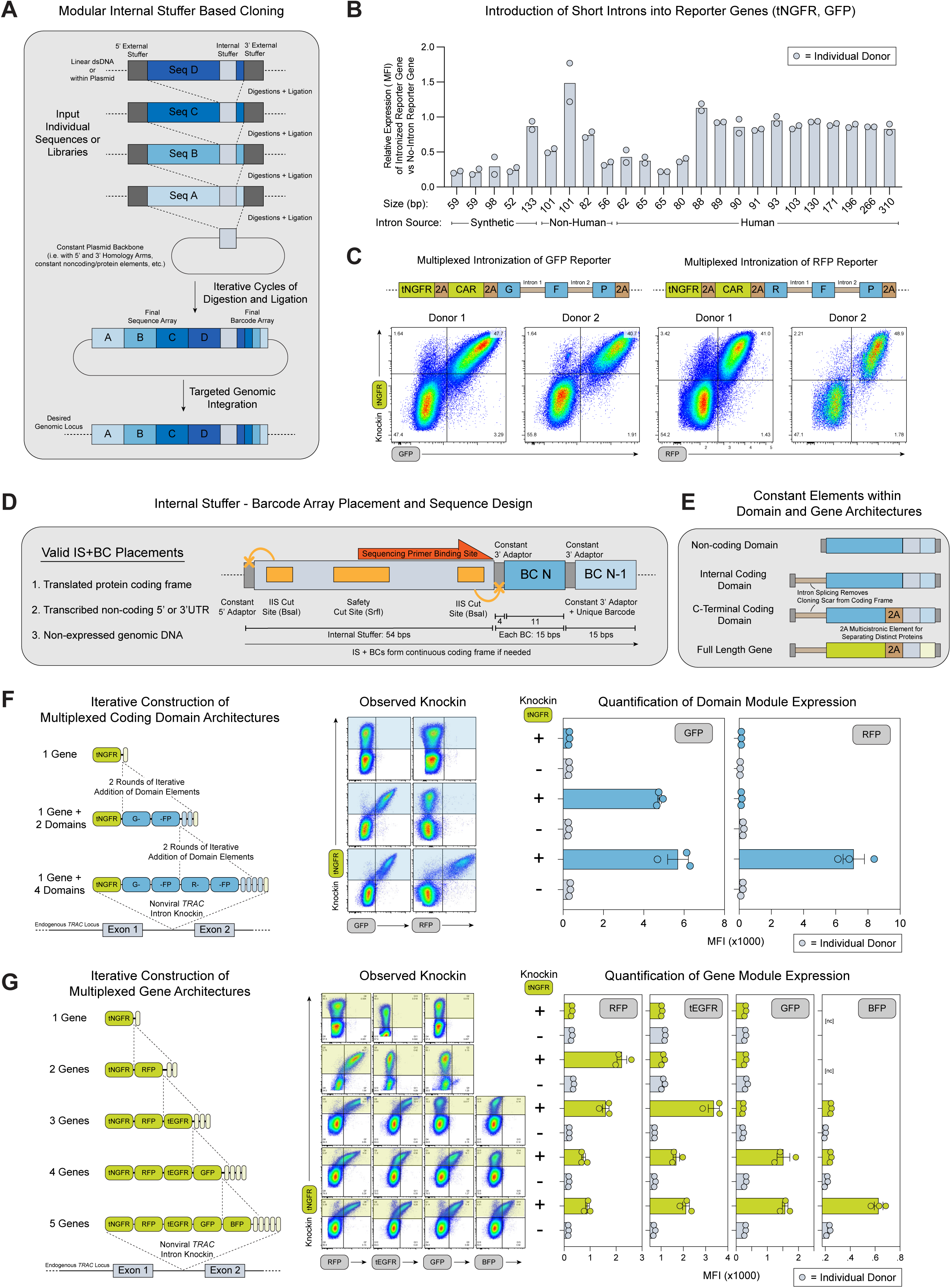
Development of Domain and Gene Architectures for Universal Pooled Knockin Screening. **(A)** Universal pooled knockin screening cloning is based on iterative cycles of digestions and ligations. Individual sequences are flanked by “External Stuffer” (ES) elements, with the functional coding or non-coding sequence component separated from a uniquely identifying barcode by an “Internal Stuffer” (IS) element. The External and Internal Stuffers contain separate Type IIS restriction enzyme sites that allow for individual sequences to serve as either an insert into a separate sequence, such as Seq B (ES Digested) → Seq A (IS digested), or as the destination of another sequence, such as Seq C (ES Digested) → Seq B (IS Digested). As the IS is preserved throughout each round, iterative cloning cycles can be continued indefinitely, with each cycle adding a new sequence to the 5’ side of the IS and a new barcode to the 3’ edge. **(B)** Relative expression of tNGFR or GFP in T cells edited with intron-containing templates, measured as MFI of knockin positive cells compared to respective reporter gene without an intron. All intron and full inthronized construct sequences provided in **Table S1**. Short synthetic or non-human introns showed variable efficiency in preserving reporter gene expression, whereas tested human introns with length above ~90 bps showed consistent expression. Protein expression measured by flow cytometry in n=2 human donors four days post editing. **(C)** Flow cytometric demonstration of multiplex intronization of GFP and mCherry reporter genes in n=2 human donors four days post editing. **(D)** Detailed schematic structure of Internal Stuffer and Barcode Array. Example DNA sequences provided in **Table S2**. **(E)** Schematic of the placement of constant sequences (introns, 2A elements) for Domain and Gene architectures. Non-coding domains do not need a preceding intron, while coding domains and genes do in order to allow for the 4bp constant cloning scar that would disrupt the coding sequence to be placed within the spliced out intronic sequence. Coding Domains that form the end of a given protein sequence and full length Genes have a 2A multicistronic element placed prior to the internal stuffer in order to separate the desired translated sequence from any additional protein products that will be subsequently added in additional cloning cycles. **(F)** Quantification of successful expression of GFP and RFP reporter genes encoded in the final Domain module architecture. GFP and RFP were separated into two Domains each, and successful expression of each was only observed when both of their domains were iterative included within the final construct. Protein expression measured by flow cytometry in n=3 human donors four days post editing. **(G)** Quantification of successful expression of RFP, tEGFR, GFP, and BFP reporter genes encoded in the final Gene module architecture. Large up to five gene multicistronic constructs did show some slight decreases in expression levels of individual genes, similar to previously reported declines in expression for highly multiplexed multicistronic cassettes^19^. nc = not collected for indicated sample. Protein expression measured by flow cytometry in n=3 human donors four days post editing.

**Supplementary Figure 2.**
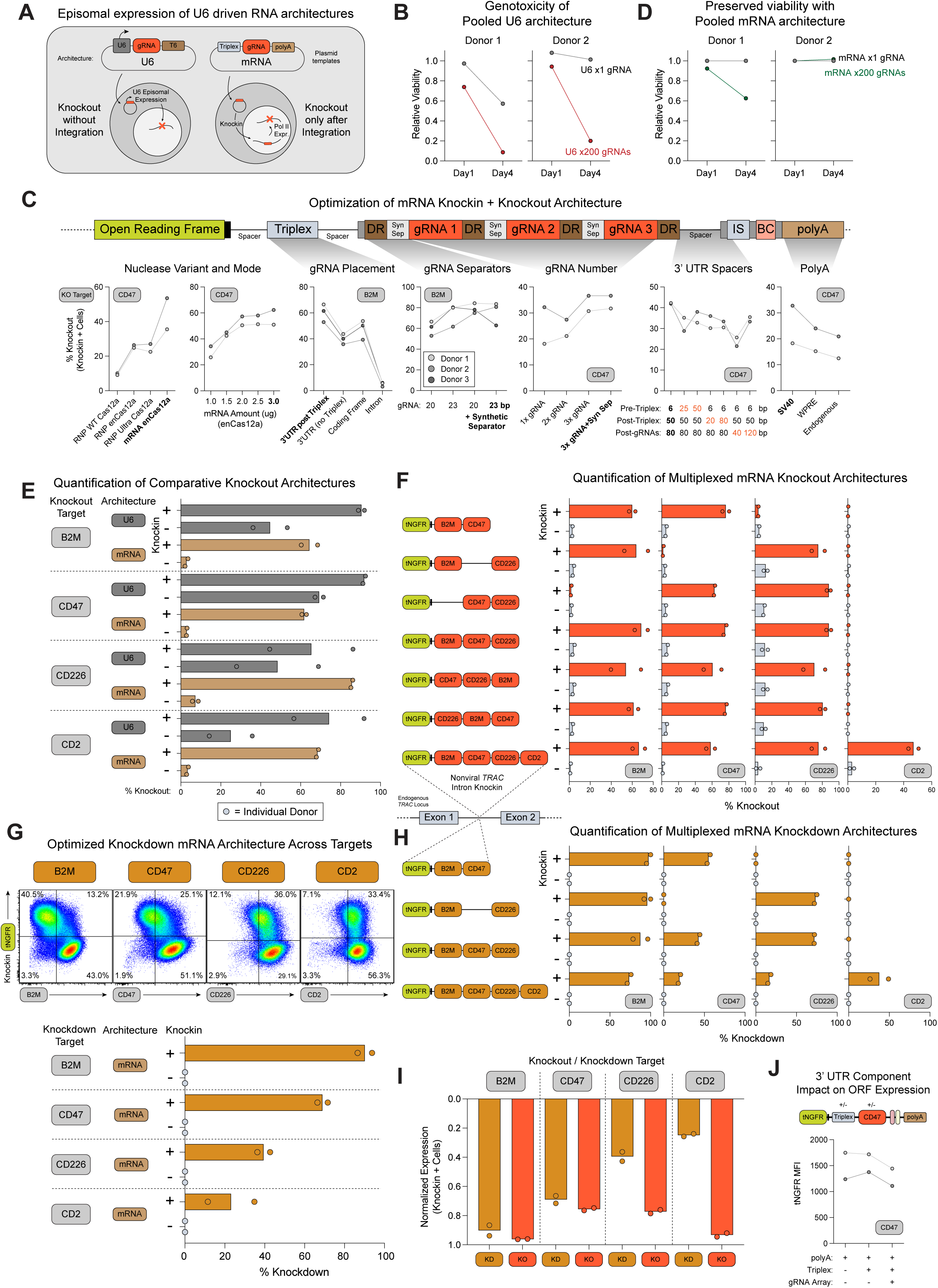
Development of Optimized mRNA Based Knockout and Knockdown Multiplexable Architectures. **(A)** Schematic of differences in timing of expression of RNA effectors between U6 driven small RNAs (such as gRNAs) compared to RNA Pol II driven mRNA based small RNA expression. U6 promoters (whether contained within a plasmid or certain viral vectors such as AAVs which express their small RNAs both episomally and after genomic integration, whereas mRNA based templates for targeted integration into an endogenous locus/endogenous mRNA transcript lack an RNA Pol II promoter within the plasmid, resulting in expression of the small RNA within an mRNA template only after successful genomic integration. **(B)** Toxicity of electroporation in human T cells of the same amount (2 ug) of DNA plasmid templates containing a U6 promoter with either a single gRNA or a small library of 200 gRNAs. Despite introducing the same total amount of DNA, the U6 driven library plasmid showed much greater toxicity (measured as relative viability to introduction of equivalent amounts of DNA plasmid without a U6 promoter), especially after four days, potentially due to the greater amounts of dsDNA break formation and resulting genotoxicity due to episomal expression of numerous gRNAs within each individual cell. **(C)** Optimization series of sequence elements and editing conditions for a mRNA based combined knockin + knockout architecture. For each given optimization series, the final chosen sequence components/design and editing conditions are bolded. Observed protein knockout in knockin positive (tNGFR+) human T cells was measured by flow cytometry four days after editing in n=2-3 donors. **(D)** Toxicity of electroporation in human T cells of the same amount (2 ug) of DNA plasmid templates containing an optimized mRNA knockin+knockout architecture with either a single gRNA or a small library of 200 gRNAs (same library as in **Figure S2B**). Unlike the observed large drop in viability for electroporation of pooled U6 driven gRNA libraries, in two human donors pooled mRNA based gRNA libraries the added viability loss from one to 200 gRNAs was small one donor and not observed in a second. **(E)** Quantification of observed protein knockout of *B2M, CD47, CD226,* and *CD2* target surface receptors using a U6 vs optimized mRNA based architectures. U6 based architectures showed knockout in both successfully edited (knockin positive, tNGFR+) and unsuccessfully edited cells (knockin negative, tNGFR-), whereas the mRNA architecture showed efficient knockout only in successfully edited cells. **(F)** Quantification of multiplexed knockout of various combinations of *B2M, CD47, CD226,* and *CD2* target surface receptors using an optimized mRNA based architecture. **(G)** Quantification of observed protein knockdown of *B2M, CD47, CD226,* and *CD2* target surface receptors using an mRNA based architecture containing a miR-E format endogenously excisable shRNA rather than a Cas12a excisable gRNA. The degree of observed knockdown varied across targets, but was selective only for successfully edited cells (knockin positive, tNGFR+). **(H)** Quantification of multiplexed knockdown of various combinations of *B2M, CD47, CD226,* and *CD2* target surface receptors using an optimized mRNA-based architecture. **(I)** Comparison of observed protein expression levels between knockout and knockdown of the same surface receptor targets. Knockouts showed higher observed reductions in protein expression in knockin positive cells compared to knockdowns. **(J)** Inclusion of a triplex and gRNA array in the 3’UTR of the integrated knockin+knockout cassette did not appear to result in major changes in the expression level of the knocked in protein component (tNGFR), as measured by tNGFR MFI. Knockin and knockout protein expression measured by flow cytometry in n=2 human donors four days post editing (**E-J**). Percent knockout calculated as the percentage of total knockin positive or negative cells within a negative protein expression gate based on unstained controls (**E,F,I**). Percent knockdown / Normalized expression calculated as the relative MFI of knockin positive or negative cells relative to stained control cells without knockdown/knockout of the target surface receptor and unstained control cells (**G-I**).

**Supplementary Figure 3.**
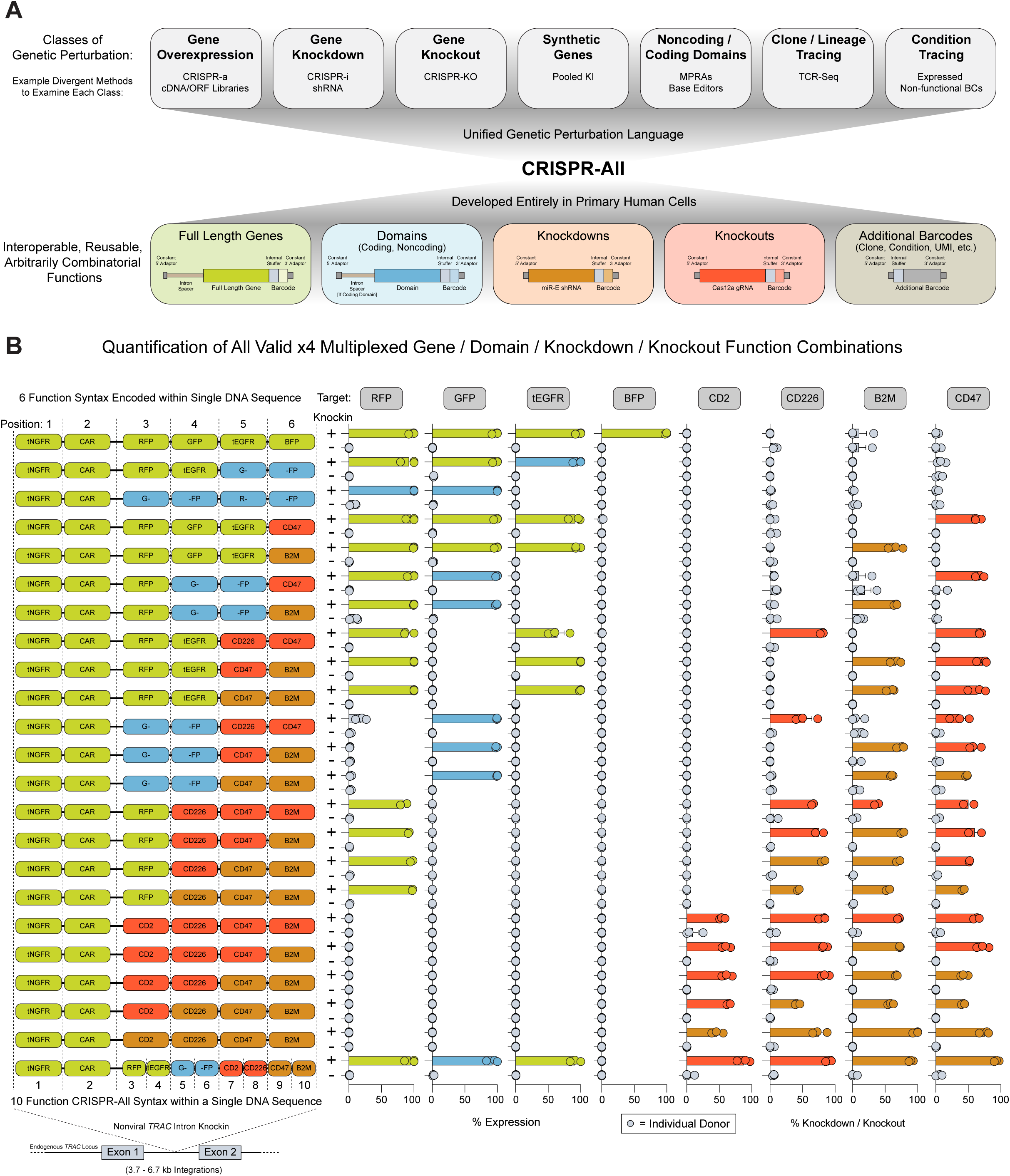
Quantification of CRISPR-All Format Multiplexed Cross-Perturbation Class Genetic Modifications. **(A)** CRISPR-All unifies bespoke, perturbation class specific methods into a single genetic perturbation language, comprised of interoperable, reusable, and arbitrarily combinatorial Gene, Domain, Knockdown, and Knockout functions. **(B)** Quantification of multiplexed gene and domain knockin and endogenous gene knockout and knockdowns across all valid x4 combinations of CRISPR-All Functions (Gene, Domain, Knockout, Knockdown), with various combinations of RFP, GFP, tEGFR, and BFP being expressed, and *B2M, CD47, CD226,* and *CD2* endogenous surface receptors being knocked out or knocked down. Each x4 perturbation function combination was knocked into the first intron of the endogenous *TRAC* locus in human T cells, along with two constant Gene functions, a tNGFR reporter and a Chimeric Antigen Receptor (for a total of x6 perturbation functions within each construct). Knockin, knockout, and knockdown protein expression measured by flow cytometry in n=3 human donors four days post editing.

**Supplementary Figure 4.**
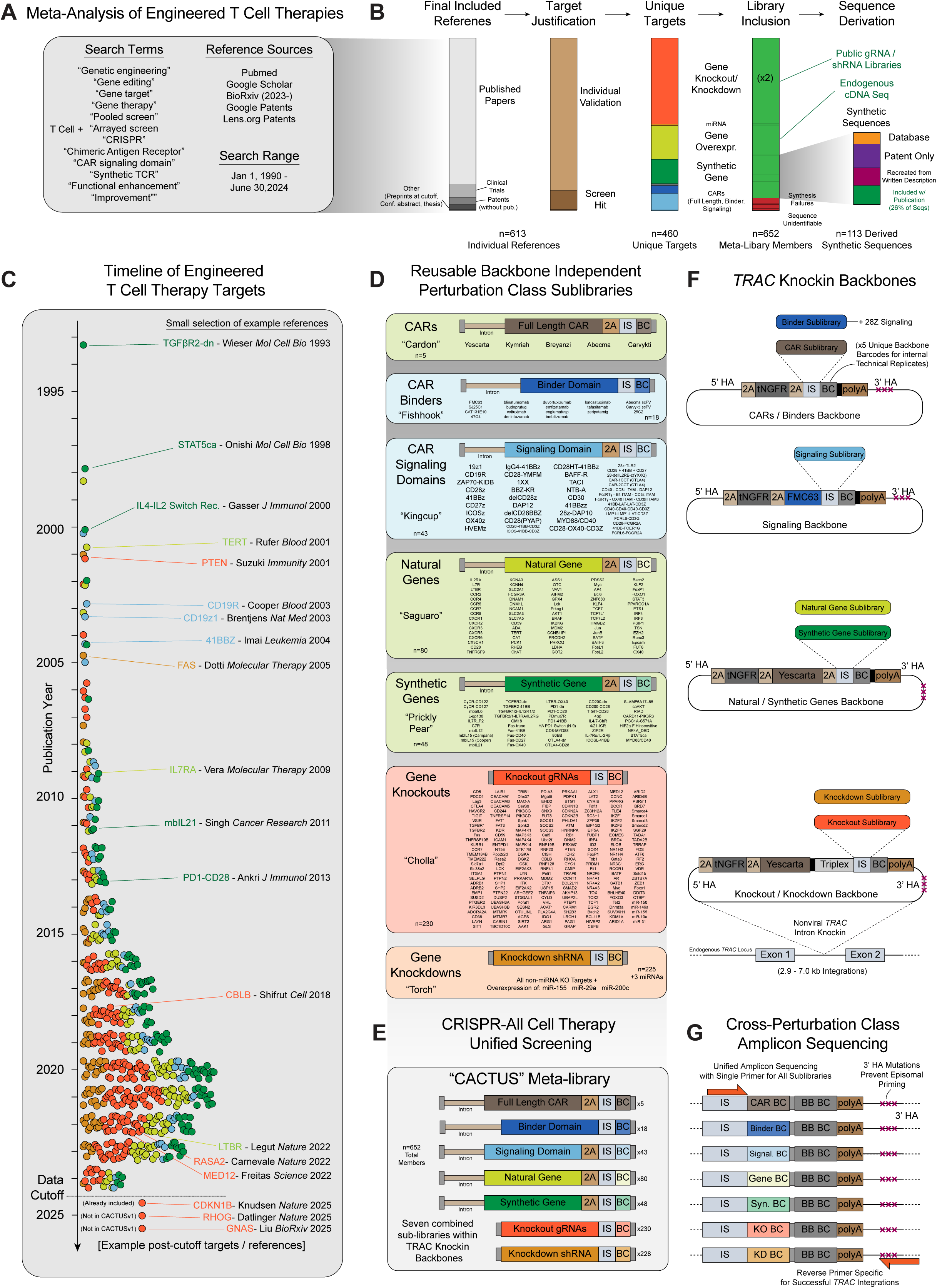
Meta-Analysis of Engineered T Cell Therapy Targets and Design of Engineered T Cell Meta-Library. **(A)** Engineered T cell therapy meta-analysis search terms, reference sources, and search range. **(B)** Meta-analysis yielded 613 individual references describing potential genetic strategies for improving some aspect of T cell function, in mouse or human T cells, with an endogenous TCR, synthetic TCR, or a CAR (**Table S10**). Specific gene or sequence targets were either nominated individually or identified as a top hit in arrayed or pooled mouse or human T cell screens (**Table S11**). Gene knockouts or knockdowns comprised the largest category of targets, with significant numbers of natural gene overexpressions, synthetic gene sequences (e.g. chimeric switch receptors, constitutively active cytokine receptors or transcription factors), and CAR signaling domains also identified. Fewer numbers of microRNAs, full length CARs (limited to FDA approved sequences), and binders (limited to CD19 binders) were identified. A final list of 460 unique genetic targets yielded a final meta-library containing 652 members (knockout or knockdown targets were included in both knockout and knockdown libraries, and a small number of sequences failed synthesis or the underlying sequence could not be identified in any publicly accessible source) (**Table S12**). The majority of published synthetic sequences (synthetic genes, full length CARs, CAR signaling domains, binders) were not included or made publicly accessible in the original publication describing them, necessitating extensive searching through public databases, patent databases, or manual recreation of sequences from a published description, in order to include them in the meta-library. **(C)** Timeline of the year of publication and perturbation class of included targets within the first version CRISPR-All Cell Therapy Universal Screening meta-library (CACTUSv1), along with a small selection of individual targets with associated reference. **(D)** Construct schematics and full list of members for each perturbation class specific sub-library included within the CACTUS meta-library. The CAR (**Table S13**), Natural Gene (**Table S16**), and Synthetic Gene (**Table S17**) sub-libraries use the CRISPR-All Gene function architecture, the CAR Binder (**Table S14**) and CAR Signaling Domain (**Table S15**) sub-libraries use the CRISPR-All Domain function architecture, the Gene Knockout sub-library (**Table S18**) uses the CRISPR-All Knockout function architecture, and the Gene Knockdown sub-library (**Table S19**) uses the CRISPR-All Knockdown function architectures. **(E)** Seven perturbation class specific sub-libraries combined together form the CRISPR-All Cell Therapy Universal Screening meta-library (CACTUSv1), all with the same standardized CRISPR-All format barcoding architecture. **(F)** Schematics of sub-library specific plasmid backbones containing constant elements along with homology arms for targeted integration into the *TRAC* locus in human T cells. Each backbone contained five separate “Backbone Barcodes” in CRISPR-All format to allow for internal technical replicates during pooled screening. **(G)** Schematic of cross-perturbation class barcode amplicon sequencing.

**Supplementary Figure 5.**
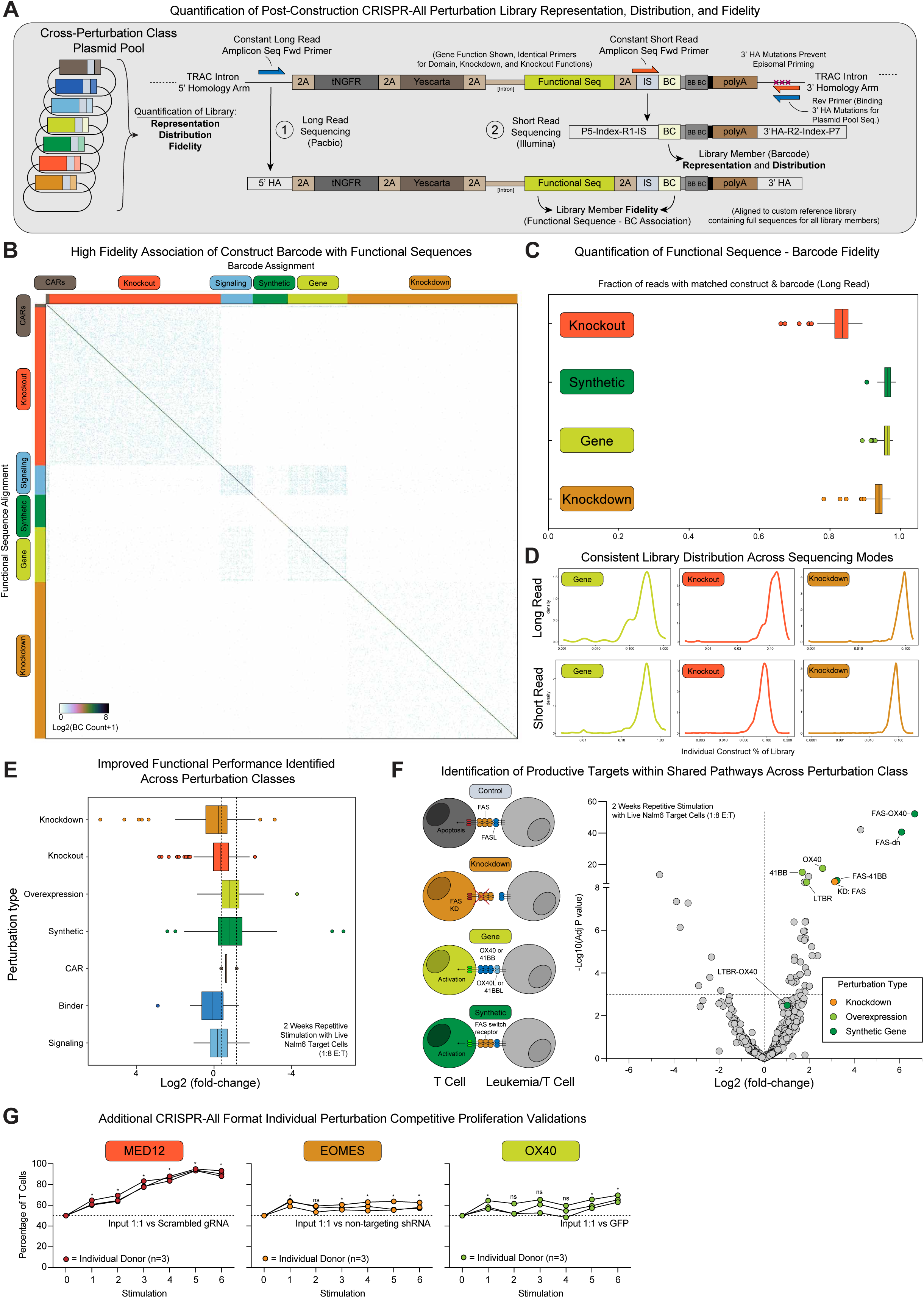
CACTUS Meta-Library Quality and Screening Validations. **(A)** Schematic for the CRISPR-All plasmid library and post-integration library quantifications of sequence fidelity (correct association of individual sequence with its designated barcode) by long read PacBio sequencing, and library member representation and distribution by short read amplicon sequencing. **(B)** High fidelity association of individual constructs with their designated barcodes by long read sequencing of the CACTUS meta-library plasmid pool. A small amount of template switching (incorrect final association of sequence to barcode) was observed, more prominently within constructs from the same perturbation class (e.g. knockout sequences more likely to template switch with other knockout sequences) (see **STAR Methods**). **(C)** Quantification by long-read sequencing of sequence to barcode association fidelity across perturbation type. **(D)** Both long read and short read quantification of CACTUS meta-library plasmid pool showed clean distributions post cloning, with over 90% of constructs with one log10 window across perturbation class specific sub-libraries. **(E)** Constructs with improved functional performance (measured as abundance after two weeks of repetitive stimulation with live Nalm6 target leukemia cells at a 1:8 E:T ratio every 2-3 days) relative to base FDA approved CARs were identified across CACTUS perturbation class specific sub-libraries, including multiple gene knockdowns, knockouts, natural gene overexpressions, synthetic genes, and CAR signaling domains. Data from n=8 human donors. **(F)** Highlighted behavior of constructs targeting similar gene pathways from across different perturbation classes that resulted in improved proliferative performance in the two week repetitive stimulation screen, in particular knockdown of *FAS*, redirection of *FAS-L* apoptotic signals using synthetic genes (e.g. FAS-OX40 switch receptor), or overexpression of natural co-stimulatory genes such as *OX40*. Data from n=8 human donors. **(G)** Individual validation of three additional CACTUS repetitive stimulation screen targets, *MED12* Knockout, *EOMES* Knockdown, and *OX40* overexpression, from different perturbation classes in a competitive repetitive stimulation assay (in comparison to indicated matched perturbation type controls) showed increased proliferation across n=3 human donors with target live Nalm6 cells (1:8 E:T). ns = not significant, * = P<0.05, Unpaired t test.

**Supplementary Figure 6.**
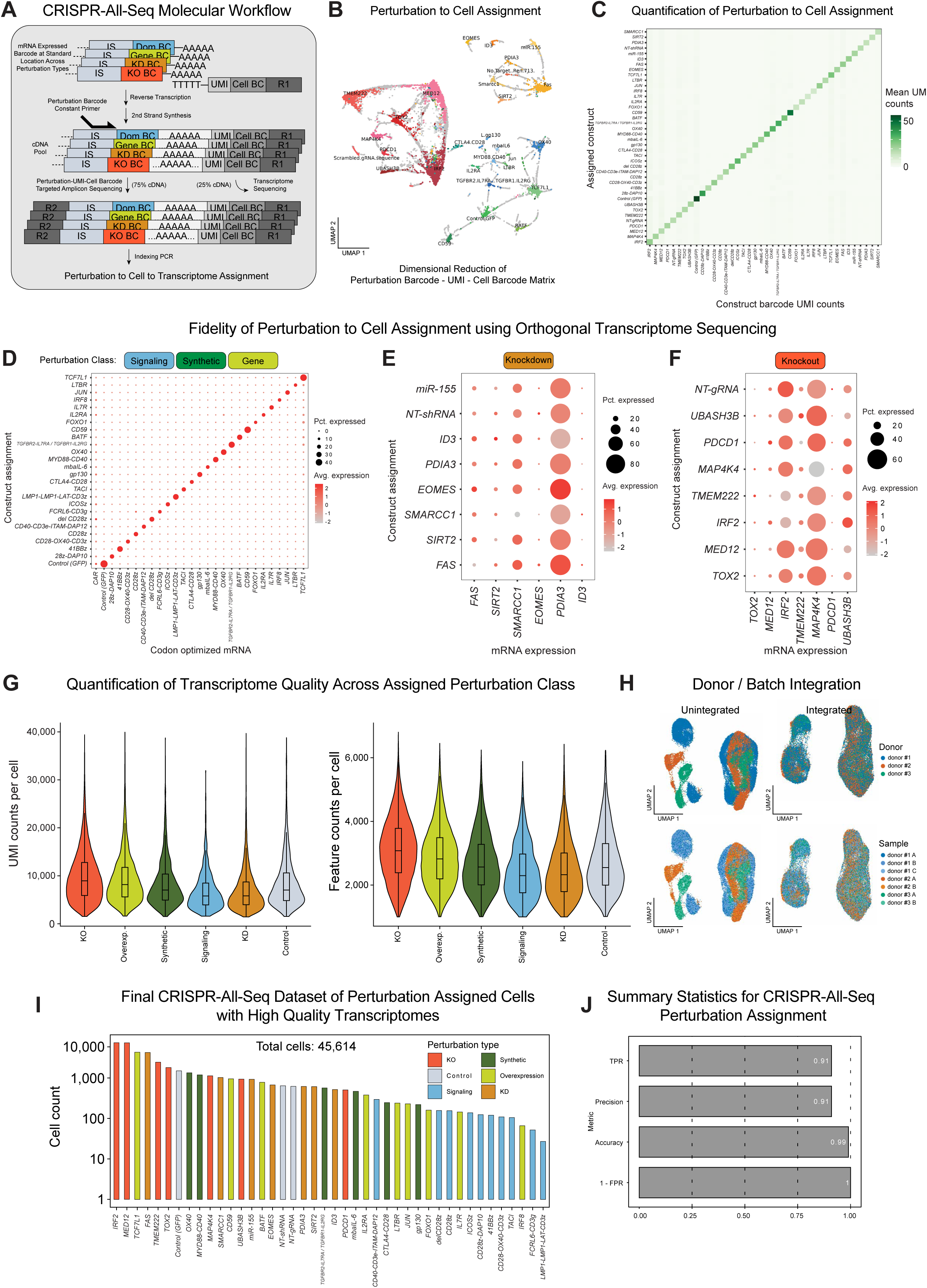
CRISPR-All-seq Assignment, Fidelity, and Transcriptome Quality. **(A)** Schematic of sequencing amplicons for associating perturbation barcodes with cell barcodes. First, mRNA is captured by pold-dT oligos with distinct cell barcodes and UMIs. Following reverse transcription and second strand synthesis, a fraction of the cDNA pool is separately amplified and indexed to generate sequencing libraries. **(B)** UMAP embedding of cell barcode by perturbation barcodes matrix where values represent UMI counts. A single representative lane from donor 1 is displayed. Knockout constructs are shown in red and pink hues, knockdown constructs in yellow hues, and gene knock-in constructs in green and blue hues. UMAP parameters are n_neighbors=15, min_dist = 0.1, spread = 1, and metric = “cosine” implemented by the ‘uwot’ R package version 0.2.3. **(C)** Heatmap display of mean UMI counts for each construct assignment. Mean UMI counts are largest for the respective assigned construct, with construct variability in ‘background’ UMI counts. We observe that constructs with greater UMI count background are typically those at greater cell abundance (see TCF7L1, FAS, MED12, or IRF2), contributing to an increased likelihood of swapping events. **(D)** Dot plot of knock-in coding sequence expression. Knock-in signaling, synthetic gene, and overexpression constructs are each codon optimized and do not align to the associated endogenous gene, when applicable. Thus, a separate alignment step using kallisto-bustools is performed to quantify expression across each construct assignment. Captured transcripts contain a CAR 5’ of the variable knock-in sequence, which is likely captured at lower abundance due to 3’ skew with 3’ scRNA sequencing. **(E, F)** Dot plots showing expression of knockdown and knockout targets for cells assigned to knockdown and knockout perturbation categories. The control shRNA and gRNA assigned cells (NT-shRNA and NT-gRNA) indicate variable baseline expression levels for knockdown and knockout targets. **(G)** UMIs and unique features per cell for each perturbation type in the CRISPR-All-seq dataset. The ‘Control’ category contains exclusively the GFP overexpression control. NT-gRNA and NT-shRNA cells are included in the ‘KO’ and ‘KD’ categories, respectively. An increase in counts for the knockout constructs is driven predominantly by the MED12-KO assigned cells which have higher feature and UMI counts and make up 33% of the knockout construct cells. **(H)** UMAP embeddings with and without integration by donor. In display column 1, cells are colored by donors and in display column 2 cells are colored by 10x lane. **(I)** Log10 scale bar plot showing the number of cells assigned to each perturbation ordered from greatest to least cells assigned. Bar colors represent the perturbation type. Cell perturbation multiplets and cells which do not have any perturbation assignment are excluded from the visualization. **(J)** Summary statistics of perturbation assignments using the subset of cells where both a perturbation barcode assignment is made, and knock-in sequence reads are detected to make sequence-based assignments. Sequence-based assignments are used as ground truth (rather than barcode-based assignments; see methods).

**Supplementary Figure 7.**
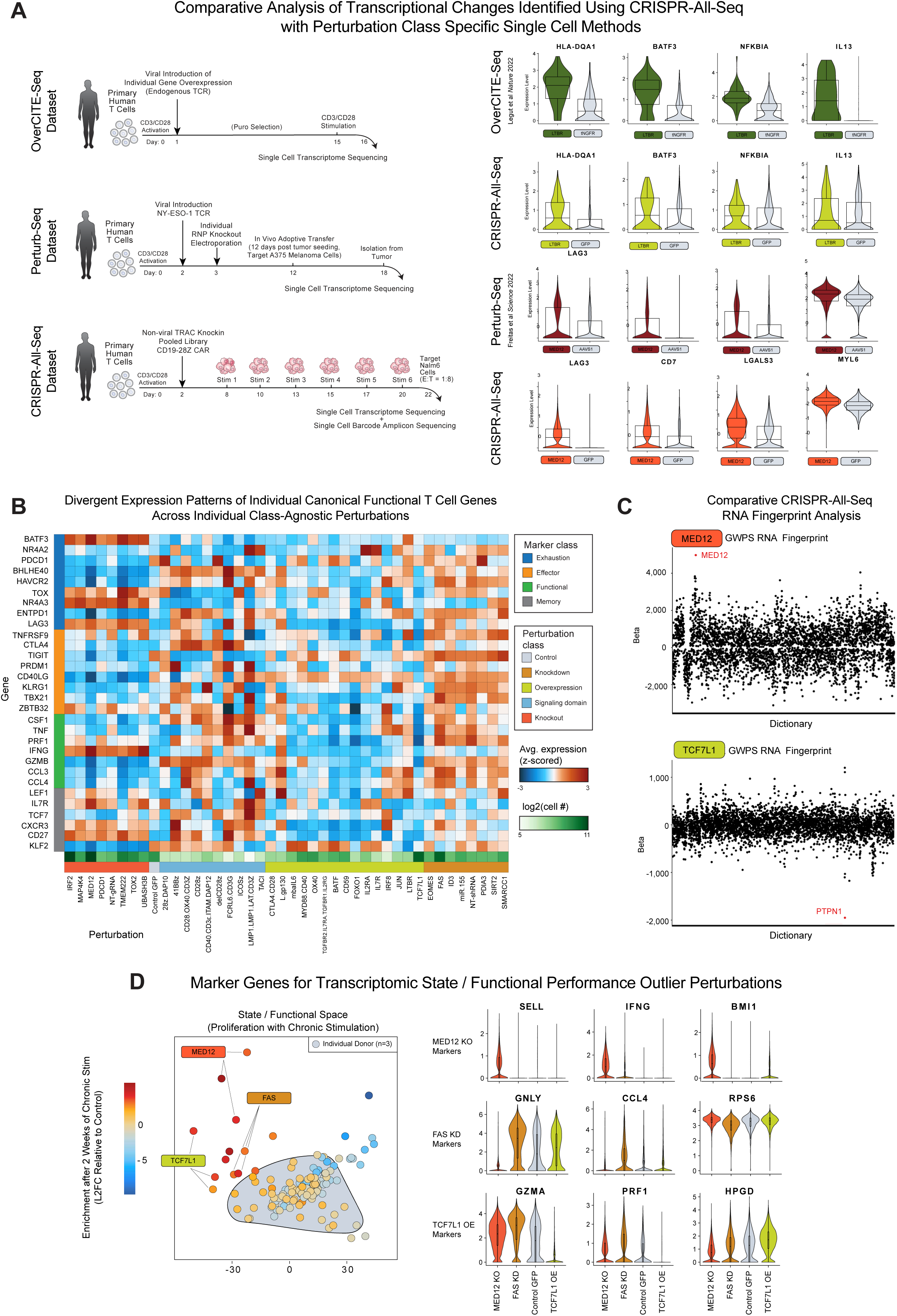
CRISPR-All-seq Analysis of Cross Perturbation Class Modified CAR T Cells after Chronic Stimulation. **(A)** Schematic of distinct experimental workflows, knock-in or transduction methods, and genetic cargo in three separate single cell experiments using primary human T cells. The first row of violin plots show expression of LTBR markers, relative to control, in the OverCITE-seq dataset. The second row shows the same markers in the CRISPR-All-seq dataset. The third row shows expression of MED12 knockout markers relative to AAVS1 knockout control in the Perturb-seq dataset. The final row displays the same markers for MED12 knockout and GFP knock-in assigned cells in the CRISPR-All-seq dataset. **(B)** Heatmap displaying z-scored expression of relevant T cell genes (rows), pseudobulked by perturbation assignments (columns). We include all cells and perturbations in this visualization but note that constructs with fewer assigned cells have more variability in gene expression z-scores. Log2 cell number is indicated per construct by the green heat bar. **(C)** RNA fingerprinting ‘Long Island City plots’ for MED12 knockout and TCF7L1 overexpression query cells using the pre-computed ‘GWPS’ (Genome-wide Perturb-seq) K562 cell reference. Bayesian regression coefficients are displayed on the y-axis. **(D)** Pseudobulk PCA representation of cells from each donor and perturbation combination. Aggregate counts are batch corrected using limma (see methods). Each donor and perturbation combination is represented by a single point. MED12 knockout, FAS knockdown, and TCF7L1 overexpression are specifically labelled with corresponding violin plots of perturbation-specific markers. The cells assigned to the control GFP knock-in category are also show for reference.

**Supplementary Figure 8.**
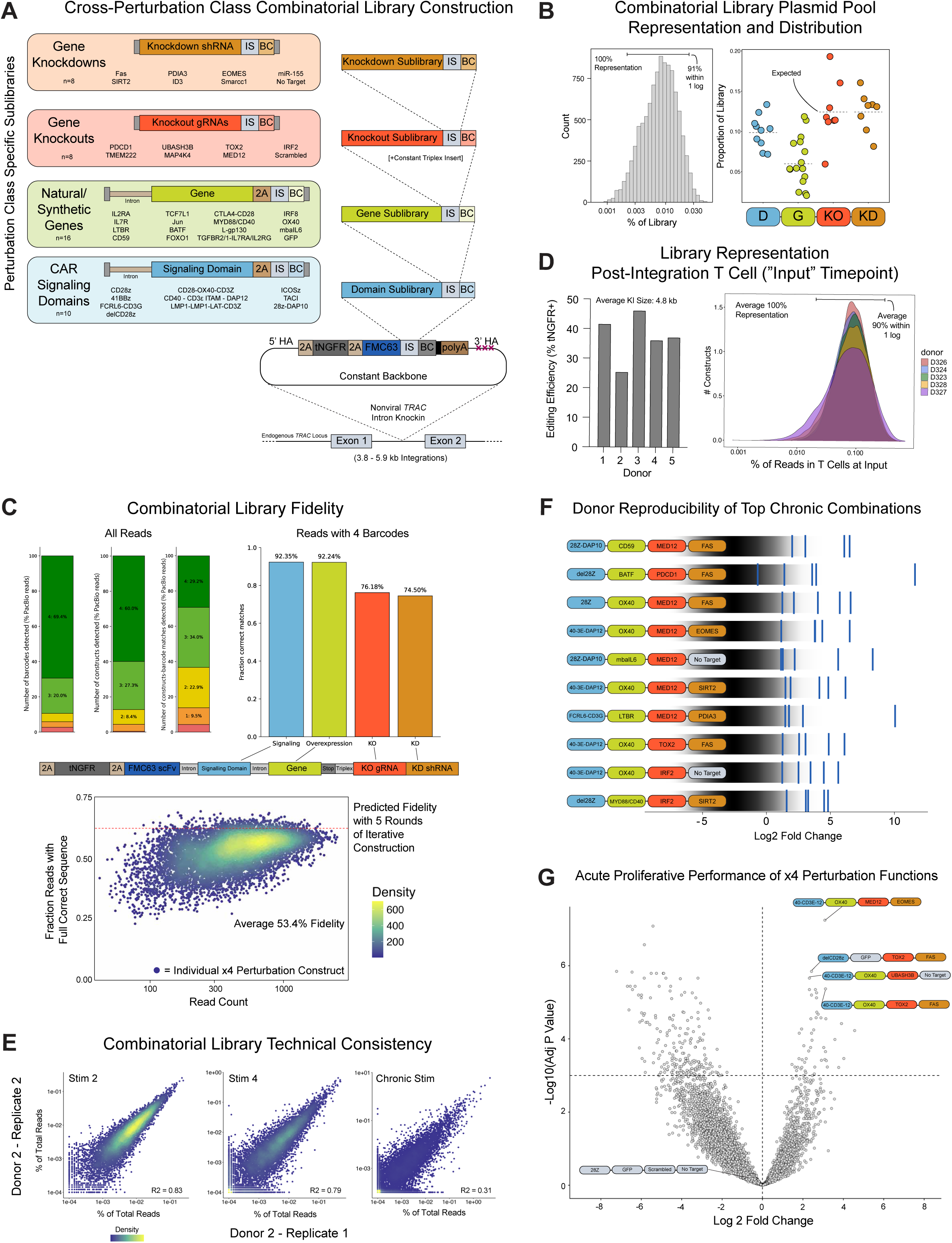
Cross-Perturbation Class Combinatorial Library Construction, Quality, and Technical Consistency. **(A)** Iterative cloning workflow of perturbation class specific sub-libraries of Gene Knockdowns, Knockouts, Natural and Synthetic Genes, and CAR Signaling Domains into a constant backbone containing a tNGFR reporter gene, an FMC63 CD19 targeting binder, and homology arms for *TRAC* integration in human T cells. **(B)** Short read sequencing of the barcodes of the combinatorial x4 perturbation function plasmid library (n=10,240 individual constructs). 100% of constructs were represented, with 91% of constructs within 1 log. Frequencies of appearance of each individual perturbation from each of the four sub-libraries also shown. **(C)** Long read sequencing of the x4 perturbation function plasmid library and computational construct and barcodes alignment enabled assessment of the fidelity of construct to barcode associations. Of all detected reads, 69.4% had 4 barcodes detected and 60.0% had all four constructs detected (constructs missing one or more components can result from incomplete digestions/ligations during the iterative cloning steps). 29.2% of all base long reads had 4 constructs, 4 barcodes and a complete match of all 4 sequences. Of reads with 4 barcodes (the only constructs sequenced and analyzed during screening), 53.4% had a complete match with all 4 construct sequences. Reads with 4 barcodes had 92.3% correct signaling Domains, 92.2% correct Genes, 76.2% correct Knockouts, and 74.5% correct Knockdowns. Given the observed degree of template switching during single perturbation function sub-library cloning during the construction of the CACTUS library (**Figure S5C**), the predicted fidelity for four perturbation function constructs was ~60%, close to the observed average fidelity of 53.4% (n=10,240 four perturbation constructs). **(D)** Editing efficiency of the combinatorial x4 perturbation function library in human T cells measured four days after electroporation in n=5 human donors. Short read sequencing of the barcode arrays from these five donors six days post editing (“Input” timepoint) showed successful high coverage library integration, with an average representation of the 10,240 constructs of 100% and an average of 90% of constructs within 1 log. **(E)** Technical consistency of the combinatorial x4 perturbation function library in technical replicates from a second human donor (Figure 6D). **(F)** Donor reproducibility across the top 10 x4 perturbation function constructs measured by average Log2 Fold Change across n=5 human donors after chronic stimulation. Each line represents an individual donor’s screen results overlayed over a grayscale distribution from the entire combinatorial library. **(G)** Results of combinatorial x4 perturbation function library abundance screening after acute stimulation in human T cells (n=5 human donors). Four days after initial stimulation with target Nalm6 cells (1:8 E:T), genomic DNA was extracted from surviving T cells and barcode arrays were amplified. Log2 Fold Change and Adjusted P-value calculated in comparison to Input timepoint.

**Supplementary Figure 9.**
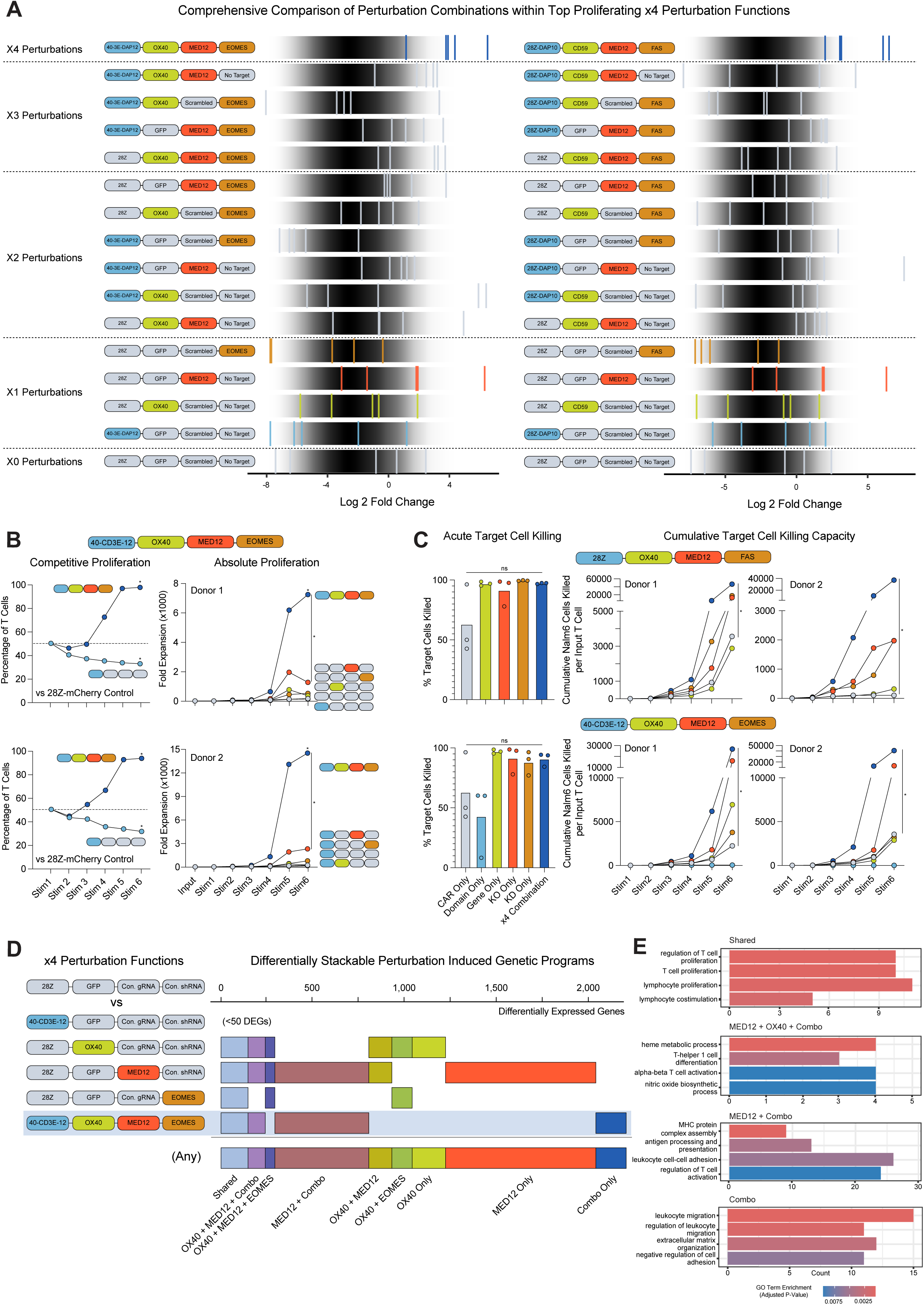
Individual Validation of a Second x4 Combinatorial Perturbation Construct. **(A)** Donor reproducibility plots for two selected top x4 perturbation function constructs along with all related x3, x2, x1, and x0 perturbation function constructs (with controls in every other position, such as GFP for Gene function and scrambled gRNA for Knockout) at the Chronic Stimulation timepoint. n=5 human donors. **(B)** Individual validation of CD40/CD3E/Dap12-OX40-MED12-EOMES x4 perturbation combination in a competitive repetitive stimulation assay (in comparison to a 28Z-mCherry-Scrambled-NoTarget control) showed increased proliferation across n=2 human donors with target live Nalm6 cells (1:8 E:T). The CD40/CD3E/Dap12 signaling domain on its own showed a mild decrease in competitive proliferation. The CD40/CD3E/Dap12-OX40-MED12-EOMES x4 perturbation combination also showed increased absolute proliferation in n=2 human donors across repetitive stimulations when individually cultured in comparison to single perturbation constructs. ns = not significant, * = P<0.05, Unpaired t test. **(C)** Acute and cumulative target cell killing assays for the 28Z-OX40-MED12-FAS and CD40/CD3E/Dap12-OX40-MED12-EOMES x4 perturbation constructs in comparison to constituent individual perturbations. In 2 of 3 donors tested at the examined 1:8 E:T ratio, the two tested combinatorial constructs, as well as OX40 individual overexpression, MED12 individual knockout, and FAS and EOMES individual knockdowns, showed increased acute killing in comparison to CAR only control T cells (Yescarta, a CD19-28Z CAR). The two tested x4 perturbation constructs also appeared to increase the cumulative number of Nalm6 target cells killed by each input T cell over the course of a two week repetitive stimulation assay in comparison to individual perturbations or CAR only control. ns = not significant, * = P<0.05, Unpaired t test. **(D)** Bulk RNA sequencing identified patterns of differentially expressed genes between CD40/CD3E/Dap12-OX40-MED12-EOMES and individual component perturbations (CD40/CD3E/Dap12, OX40, MED12, FAS) relative to control edited T cells (28Z-GFP-Scrambled-NoTarget) after two weeks of chronic stimulation. n=3 human donors. All individual or overlapping sets of >50 DEGs shown in n=3 human donors. **(E)** Gene Ontology term enrichment in selected overlapping sets of differentially expressed genes which included the CD40/CD3E/Dap12-OX40-MED12-EOMES x4 perturbation construct (“combo”).

## METHODS

### Cell Lines

Nalm6 cells were obtained from American Type Culture Collection (ATCC) and maintained in RPMI (Fisher Scientific) supplemented with 10% FBS. The Nalm6 line used for all experiments was genetically modified by targeted non-viral gene editing to insert an SFFV promoter immediately before the start codon of the endogenous *BCMA* start codon, followed by sorting of BCMA+ Nalm6 cells to generate a pure target cell line expressing both CD19 and BCMA. Nalm6 cells were cultured for up to ~20 passages before discarding cultures are returning to low passage number frozen aliquots of initial edited and purified cell line stocks.

### Primary Cells

PBMCs from healthy human blood donors were collected under an approved IRB protocol by the Stanford Blood Center and used to isolate human T cells.

### Primary Human T Cell Isolation and Culture

Leukoreduction chambers (LRS) from processing of platelet donations were used to isolate PBMCs using density centrifugation with Ficoll (Lymphoprep, StemCell) within SepMate tubes (StemCell) according to manufacturer’s instructions. Next, primary human CD3 positive T cells were isolated by negative selection using Human CD3 T Cell Enrichment kit (StemCell) according to manufacturer’s instructions. Isolated primary human CD3 T cells were counted using an automated cell counter (Countess, Thermo) and activated using anti-human CD3/CD28 dynabeads (Cell Therapy Systems, Thermo) at a 1:1 ratio in XVivo 15 media (Lonza) supplemented with 5% FBS (MilliporeSigma) and 50 U/mL of human IL-2 (Peprotech). T cells were activated at 1:1 ratio of cells to dynabeads and initially cultured in standard tissue culture incubators at approximately 1e6 cells / mL media. After gene editing/electroporation, T cells were counted and reseeded at approximately 1e6 cells / mL XVivo 15 media with fresh IL-2 every 2-3 days.

### Non-viral Gene Knockins

Two days after activation, human T cells were harvested, dynabeads were magnetically removed by incubating for two minutes at room temperature on a magnet (EasySep Magnet, StemCell), and cells were counted using an automated cytometer. For electroporations, one to two million T cells per editing condition were gently pelleted by centrifugation at 90G for 10 minutes, followed by careful aspiration of the supernatant. T cell pellets were resuspended in 20 uL per editing condition in P3 Buffer (Lonza). Unless indicated in figure legends, all editing was performed using an enCas12 mRNA template, along with plasmid HDR templates and a plasmid expressed Cas12a gRNA targeting the first intron on the human *TRAC* locus (5’ GCAGACAGGGAGAAATAAGGA 3’). enCas12a mRNA was produced by in vitro transcription (NEB HiScribe® T7 High Yield RNA Synthesis Kit) according to manufacturer’s instructions, with provided uridine replaced by N1-Methylpseudouridine-5’-Triphosphate (TriLink), purified by SPRI cleanup, and normalized in H2O to 1 ug/uL. For electroporation, 3.0 uL of enCas12a mRNA was mixed with 0.5 ug of plasmid HDR template at 1 ug/ug, 0.5 ug of a gRNA expression plasmid at 1 ug/uL (~4kb in size, with a U6 promoter driving expression of the *TRAC* intron knockin Cas12a gRNA), and 1.5 ug of small “helper” plasmid at 1 ug/uL (also ~4kb in size). For every editing condition, 5 uL of mRNA / plasmid mixture was then mixed with primary T cells resuspended in 20 uL P3 buffer and 25 uL total were electropoated per well on a Gen2 Lonza 4D electroporation/nucleofection system using 96 well plate attachment and 96 well plate cuvettes, using pulse code EO-151. Immediately following electroporation, 75 uL of pre-warmed XVivo 15 media was added to each cuvette, and cells were rested within the cuvettes for 15 minutes in a standard 37C Tissue Culture Incubator prior to moving to culture plates or flasks.

### Individual DNA Constructs and HDR Templates

All individual non-CRISPR-All format DNA constructs used in the study were generated using standard cloning methods, primarily using PCRs (Q5 ultra II HotStart Polymerase, NEB), Gibson Assemblies (NEBuilder HiFi DNA Assembly Master Mix, NEB) and bacterial transformations (Mix & Go! Competent Cells - DH5 Alpha, Zymo). DNA constructs were sequence confirmed by Sanger Sequencing (Elim Bio) or whole plasmid next-generation sequencing (Plasmidsaurus). For gene editing, plasmid Homology Directed Repair Templates were produced by bacterial culture and standard plasmid preparation (Zymo Midiprep and Maxiprep kits) according to manufacturer instructions, including endotoxin removal steps. Final plasmid HDR templates were eluted in molecular grade water, quantified (Nanodrop), and diluted to final concentrations of 1 ug/uL for electroporations. Annotated genebank files for all DNA constructs (described in **Tables S1-S3, S9-15, S18-19)** used in the study are available upon request.

### CRISPR-All Construct Design

CRISPR-All format architectures for Domain, Gene, Knockout, and Knockdown functions are described in **Figures 1**, **2**, **S1**, **S2**, and **S3**, and detailed sequence examples are annotated in **Table S2.** Briefly, each functional sequence (cDNA sequence, coding domain, gRNA, etc.) is separated from a unique 11 bp barcode by an internal stuffer element. Each Functional Sequence-Internal Stuffer-Barcode cassette is flanked by an External Stuffer on both the 5’ and 3’ ends. A Functional Sequence-Internal Stuffer-Barcode cassette can be excised from a plasmid backbone (or PCR amplicon in the case of sequences short enough to be synthesized as ssDNA oligos, such as Knockdown and Knockout functions) by Type IIS restriction digestion sites in the External Stuffers, always leaving a constant “GTAA” four bp overhang on the 5’ end and a constant “AGCG” four bp overhang on the 3’ end, and allowing the cassette to serve as an insert into any other cassette containing a “opened” or digested Internal Stuffer. A Functional Sequence-Internal Stuffer-Barcode cassette can serve as the destination for another cassette by digesting open its Internal Stuffer again using Type IIS restriction digestion sites contained within the Internal Stuffer, similarly always leaving a constant “GTAA” four bp overhang on the 5’ end and a constant “AGCG” four bp overhang on the 3’ end. All barcodes were selected from a whitelist whereby each 11 bp barcodes has a hamming distance of at least 3 from any other 11 bp barcode. The Internal Stuffer contained a constant primer binding sequence (5’ CAACAATGTGCGGACGGCGTTG 3’) to allow any CRISPR-All format barcode array to be amplified with a constant sequence, from genomic DNA or from mRNA converted to cDNA such as during CRISPR-All-seq single cell workflows. For natural or synthetic gene overexpression and domains, the integrated coding sequence was codon optimized with two primary objectives. First, sequences were human codon optimized (TWIST API or custom script) to enable gene synthesis which requires minimizing repetitive stretches and DNA homopolymers while maintaining codon usage rates which mirror the human genome. Second, we aim to minimize the likelihood of inadvertent splicing within the coding sequence by scoring each optimized sequence and swapping in synonymous codons at predicted splice sites^118^. CRISPR-All format genome-wide knockout and knockdown library sequences are provided for the human genome (**Tables S4-5**). Final sequences for CRISPR-All Gene and Domain Functions were largely synthesized as clonal genes with NGS sequence verification (TWIST, Thermo), while CRISPR-All Knockout and Knockdown Functions were largely synthesized as gene fragments for individual constructs (TWIST, IDT) or as ssDNA oligos for pools (TWIST, IDT).

### CRISPR-All Construct Cloning

Internal Stuffer digestions were performed with BsaI (NEB) with safety/cloning efficiency secondary digestions by SrfI (NEB). External Stuffer digestions were performed with BbsI (NEB) with safety/cloning efficiency secondary digestions by PmeI (NEB). All digestions were performed with 1 ug of plasmid or PCR template with 2.5 uL of each indicated enzyme with manufacturer recommended buffer (NEB Cutsmart) in 50 uL total volume for 2+ hours at 37C on a thermocycler (Bio-Rad T100). Digested inserts and backbones were ligated using the ssDNA nick/sticky end specific ligase T7 (NEB) according to manufacturer’s protocol, except for a 10 fold reduction in total DNA amount, which was found to significantly reduce the incidence of ligated concatemers (which can lead to barcode template switching). Where possible, final destination plasmid backbones contained ampicillin resistance and CRISPR-All format inserts were synthesized/constructed into cloning vectors containing kanamycin resistance. For pooled libraries, a SPRI purification was performed after both digestion and ligation steps. For individual CRISPR-All formal plasmids, subsequent chemical transformation and HDRT purifications were performed as described above. For CRISPR-All pooled libraries, up to 100 ng of purified ligation product was electroporated into Endura electrocompetent cells (Lucigen) using a Bio-Rad Gene Pulser XCell instrument, recovered for 1 hour at 37°C with shaking and then grown for 20-28 hours at 30°C prior to plasmid preparation using ZymoPURE II kits (Zymo Research) with associated endotoxin removal steps and eluted in H2O. The sequences of a sample of individual clones were confirmed by whole plasmid next-generation sequencing (Plasmidsaurus) and the barcode architecture of the plasmid pool confirmed by Sanger Sequencing (Elim).

### CRISPR-All Cell Therapy Universal Screening (CACTUS) Meta-Library Construction

A literature meta-analysis of genetic targets for T cell functional engineering was conducted using indicated terms and search strategy (**Figure S4A**) with inclusion cutoff dates of January 1, 1990 – June 30, 2024. Included references nominating individual targets and pooled screens nominating top hits are described in **Tables S7** and **S7**. The final list of all included targets is described in **Table S8**. The Full-Length CAR (“Cardon”, **Table S9**), Binder Domain (“Fishhook”, **Table S10**), CAR Signaling Domain (“Kingcup”, **Table S11**), Natural Gene Overexpression (“Saguaro”, **Table S12**), and Synthetic Gene (“Prickly Pear”, **Table S13**) sub-libraries were each generated by clonal gene synthesis of all sub-library members, mixed equimolar, and then cloned in a pool into indicated plasmid backbones (**Figure S4F**) using the CRISPR-All cloning workflow described above. The Gene Knockout (“Cholla”, **Table S14)** and Gene Knockdown (“Torch”, **Table S15**) sub-libraries were ordered as pooled ssDNA oligos, amplified by PCR for 10 cycles with primers binding to the 5’ and 3’ External Stuffers to convert to linear dsDNA, and then digested and ligated into indicated plasmid backbones as described above. The Binder Domain sub-library had an additional constant CRISPR-All format 28ζ signaling domain introduced in a subsequent cloning round to create a complete CAR sequence.

### CRISPR-All Barcode Fidelity Assessment

In order to adequately sample and quantify the proper association of constructs with barcodes, PacBio HiFi sequencing was performed on PCR products generated using the below primers:

Forward: 5’-GGAACCGGTGCTGGAAGTGGT-3’

Reverse: 5’-CACTGAGCCTCCACCTAGCCT-3’

The ‘base’ library (Figure 4) and the combo library (Figure 6) were each separately sequenced targeting 1,000 reads per unique construct. For the base and combo libraries barcodes were identified via brute force search using a custom python script (see code availability). To minimize possible off-target barcode detections, a constant “AGCG” cloning scar is concatenated to each end of barcodes resulting in a 19 basepair sequence (4-bp scar, 11-bp barcode, 4-bp scar). Each knockdown sequence is a 97-bp sequence with a 22-bp siRNA sequence. Each contains a constant 5’ sequence, stem loop, and 3’ sequence. Each Cas12 knockout sequence contains a three guide array each targeting the same gene with constant sequences upstream, downstream, and in between each gRNA sequence. Gene knock-in sequences and CAR signaling domain sequences contain exclusively the coding sequence. All alignment steps were performed using minimap2 version 2.28-r1209 with the ‘map-hifi’ preset^119^.

For the base library, due to amplicons spanning the full transcript which contains a CD19 CAR, the FMC63 binder domain is overrepresented as the primary alignment because it matches the CAR binder, despite the read containing a different construct sequence. Thus, the FMC63 binder domain and barcode were excluded from the final barcode and sequence references. SAM file outputs were sorted and indexed with samtools version 1.21 and alignments for each read were extracted with a custom python script. Detected barcodes and construct alignments were merged into a single file and correctness values are determined by dividing the number of reads with matching construct and barcode assignments by the total number of reads. In the heatmap display in Supplemental Figure 5B, all constructs where a binder barcode is detected are excluded from visualization. Similar to the reason for exclusion of the FMC63 binder, we observed that a majority of reads where a binder barcode is detected coincide with a CD28ζ signaling domain as the assigned construct. This signaling domain matches the signaling domain sequence found in the CD19 CAR, and was filtered because it occluded our ability to directly measure barcode to sequence fidelity.

For the combo library, four separate construct references were built to reflect the four types of perturbations encoded in each position. After barcode detection and construct alignment, four barcode and four construct vectors are generated for each read. The separately computed fidelities for gene knock-in, signaling domains, gene knockout, and gene knockdowns are computed by filtering to only the reads where a barcode and construct are detected and dividing the number of matches by the total number of remaining reads. We observe heightened barcode swapping for the gene knockout and gene knockdown constructs, which share greater homology than the gene knock-ins and signaling domains and were amplified as pools prior to cloning. Of note, the amplification step to prepare the long read sequencing libraries may result in additional barcode swapping events and a minor underestimation of true barcode fidelity.

### CRISPR-All Barcode Amplicon Sequencing

For all amplicon sequencing screens, sequencing libraries were generated by amplicon sequencing using two rounds of PCRs with NEB Next Ultra II Q5 polymerase and primers purchased from IDT. The first PCR (98°C for 30”; 18 cycles of 98°C for 10”, 70°C for 30[s, 72°C for 45”; 72 °C for 2’) used a forward primer annealing to the “Internal Stuffer” with an Illumina Read 1 overhang and a reverse primer annealing to a region of Intron 1 of the *TRAC* locus (corresponding to the location of “Homology Arm Mismatches” with the original HDR Template plasmids) with an Illumina Read 2 overhang. Following SPRI purification at a 1X volume ratio and input normalization, PCR (98°C for 30”; 12 cycles of 98°C for 10”, 60°C for 20”, 72°C for 25”; 72 °C for 2’) to append indexes and Illumina sequencing adaptors was performed using standard Illumina Nextera primers. SPRI purified, normalized and pooled libraries were sequenced on a NextSeq instrument (Illumina).

### CRISPR-All Barcode Abundance Analysis

Each CRISPR-All format construct contains a unique barcode array, with each barcode having a minimum hamming distance of three with any other barcode. Reference tables for barcode arrays in all described pooled screens are available in **Tables S16, S18,** and **S19**. In R, a reference barcode table was stored as a PDict (Biostrings package) object, raw fastq files were read in using readFastq (ShortRead package) and abundance was calculated using vwhichPDict (Biostrings package), with no mismatches. Due to a drop in sequencing quality in regions of constant sequence “AGCG” cloning scars were trimmed from fastqs using a custom bioawk script^120^. Log2FC was calculated by comparing barcode abundance between indicated conditions (e.g. “Chronic Stimulation” timepoint in comparison to “Input” timepoint) using DESeq2 with default settings (design = ~ Donor + Condition; where “Donor” indicates the human donor and “Condition” indicates the populations compared). Raw barcode counts are available for all screens in **Tables S17** and **S20**.

### Flow Cytometry

All flow cytometry data shown was acquired using a Bio-Rad ZE5 analyzer. All antibodies used for flow cytometric experiments and staining dilutions are described previously^19^. Briefly, for flow cytometric staining, samples of approximately 100,000 T cells per analyzed condition were placed into round bottom 96 well plates and centrifuged for 5 minutes at 300G. After discarding the supernatant, each well was resuspended in 20 uL of FACS Buffer (PBS + 2% FBS) mixed with desired antibodies at indicated dilutions, and incubated at 4C for 20 minutes in the dark. Cells were washed once with FACS Buffer and resuspended in FACS Buffer for analysis. Flow cytometric data was analyzed using FlowJo v10 software.

### *In Vitro* Repetitive Stimulation Pooled Screening Assay

Starting on day 6 post T cell editing (Day 8 post initial T cell activation), CAR T cells were co-cultured in X-Vivo15 Media with 50 U IL-2/mL with Nalm6 cells at effector to target (E:T) ratios indicated in figure legends at a density of ~0.5e6 cells/cm2 and ~1e6 cells/ml. Every two to three days (Day 10, 13, etc. post initial T cell activation), the co-culture(s) was sampled (50 uL in duplicate/triplicate) and stained with antibodies against NGFR (CAR) and CD19 (Nalm6) and a viability dye. Samples were acquired with CountBright Plus Absolute Counting Beads (Thermo Fisher) to determine the counts of CAR T cells and Nalm6 cells. The co-culture was re-seeded to the predefined E:T. For pooled screens a minimum coverage of 1000X (at least 1000 edited T cells on average per construct in the library) was maintained at initial and every reseeding timepoint.

### *In Vitro* Competitive and Absolute Proliferation Assays

The “Repetitive Stimulation Assay” was performed with a 1:1 mix of T cell populations: one population edited with a CRISPR-All format construct and the second edited with a corresponding control CRISPR-All format version (e.g. containing a GFP control gene instead of a natural gene overexpression). When assessing the co-cultures by flow cytometry for re-seeding, the CAR (NGFR+) population was sub-gated based on GFP status to determine the proportion of each T cell population. For absolute proliferation assays, no control cells were mixed and the examined CRISPR-All format construct was initially seeded and reseeded on its own back to indicated E:T ratio (generally 1:8 T cells to Nalm6) with the degree of T cell proliferation and target cell killing determined by flow cytometry with CountBright Plus Absolute Counting Beads (Thermo Fisher).

### *In Vitro* Cancer Target Cell Killing Assays

At eight days post non-viral gene editing, edited CAR T cells (either bulk populations without selection or selected cells as described) were mixed at indicated E:T ratios with target Nalm6 leukemia cells in 96 well plates, with 40,000 Nalm6 cells and varying numbers of T cells per well. Cell killing was assessed by flow cytometry at 48 hours, and the percentage of Nalm6 tumor cell killing was calculated by taking 1 – (# Nalm6 cells alive in experimental condition / # Nalm6 cells alive in no T cell conditions).

### CRISPR-All-seq Library Preparation

CRISPR-All-seq single cell sequencing was conducted using GEM-X Universal 3’ Gene Expression v3 reagents (10X). As CRISPR-All format barcodes are present in the 3’UTR of an expressed endogenous transcript, live CAR T cells (NGFR+) were sorted from each donor after chronic stimulation. Sorted cells were counted (Countess 3) and 25,000 cells per lane were loaded onto a Chromium Controller (10X) across seven lanes. The first donor accounts for three lanes and donors two and three are each loaded separately on the remaining four lanes. Following reverse transcription, second strand synthesis, and initial cDNA library amplification (~10 cycles) per manufacturer instructions (10X), the cDNA library was split, with 25% of the cDNA library being used for gene expression library generation and sequencing (thus linking single cell barcodes to gene expression values), and 75% of the cDNA library was used for targeted amplicon sequencing of the CRISPR-All format barcodes (thus linking single cell barcodes to the CRISPR-All perturbation barcode). Gene expression libraries were prepared according to manufacturer’s instructions, quantified using an Agilent Bioanalyzer and sequenced on a NovaSeqX instrument with the following parameters: 28 bp Read 1, 10 bp i7 Index, 10 bp i5 Index, 91 bp Read 2.

cDNA library CRISPR-All barcode amplicon sequencing largely followed the amplicon sequencing protocol for genomic DNA amplifications. Briefly, cDNA CRISPR-All barcode libraries were generated by amplicon sequencing using two rounds of PCRs with NEB Next Ultra II Q5 polymerase and primers purchased from IDT. The first PCR (98°C for 30”; 18 cycles of 98°C for 10”, 72°C for 45”; 72 °C for 2’) using a forward primer annealing to the “Internal Stuffer” with an Illumina Read 2 overhang and a reverse primer annealing to the TruSeq Read 1 constant sequence (**Figure S6A**). Following SPRI purification at a 1X volume ratio and input normalization, a second PCR (98°C for 30”; 12 cycles of 98°C for 10”, 60°C for 20”, 72°C for 25”; 72 °C for 2’) to append indexes and Illumina sequencing adaptors was performed using a standard Illumina Nextera Read 2 binding primer and a TruSeq Read 1 binding primer. The final cDNA derived CRISPR-All barcode library was SPRI purified, normalized, and pooled libraries were sequenced on a NextSeq550 instrument (Illumina).

### CRISPR-All-seq Analysis

Cellranger version 6.0.0 was used to process fastq files sequenced on a NovaSeqX targeting 20,000 reads per cell^121^. Filtered gene expression matrices were loaded in R version 4.5.0 with Seurat and SeuratObject version 5.2.0^122^.

In order to assign knock-in construct barcodes with individual cells, amplicon sequencing reads were divided by read index position into cell barcode (R1), UMI (R1), and construct barcode (R2). An unfiltered cell barcode by construct barcode matrix of UMI counts was generated and filtered to include only called cell barcodes using a custom script. No barcode sequence correction or UMI collapsing to correct for sequencing errors was performed. deMULTIplex2^123^ was used with default parameters to assign cells to constructs. Each cell is assigned to a specific construct category, multiplet, or the negative category. We exclude multiplet-assigned cells from analysis because they are due to bi-allelic knock-in or cell doublets which we are unable to confidently distinguish.

A donor-integrated dimension reduction was generated using the IntegrateLayers function in Seurat using Harmony by setting the method equal to HarmonyIntegration^124^. Unless otherwise specified, perturbation markers were identified using FindMarkers with default parameters which uses a Wilcoxon rank sum test efficiently implemented by presto^125^.

Pseudobulk matrices were generated by collapsing cells on unique donor and construct combinations using the AggregateExpression function in Seurat. The resulting matrix was log normalized and batch effects were removed with the remoeBatchEffects function from limma version 3.63.13^126^ with the donor and perturbation type as batch variables. Log2 fold-change values were computed using the number of GFP-assigned cells as a baseline.

### CRISPR-All-seq Comparison with LTBR OverCITE-seq and MED12 Perturb-seq signatures

Single cell matrices from the literature were downloaded from GEO^127^ under GEO accession number GSE193736 for the OverCITE-seq data and GSM6568647 and GSM6568648 for the Perturb-seq data.

We determined cell overexpression assignments for the OverCITE-seq dataset using the open-reading frame counts matrix and found cell assignments using the same deMULTIplex2 procedure described in the previous methods section. The gene expression matrix was then normalized using NormalizeData in Seurat with default parameters. An LTBR gene expression signature was determined using markers from the CRISPR-All-seq object. LTBR-assigned cells were compared against GFP-assigned cells using a logistic regression test regressing for donor effects and setting ‘min.pct’ to 0.02. The top 50 differentially expressed genes were selected, excluding LTBR itself, for computing perturbation signatures. The signature was then added to the OverCITE-seq and CRISPR-All-seq objects using the AddModuleScore function in Seurat.

We separately downloaded gene expression matrices for MED12 knockout cells and AAVS1 knockout cells (control) and merged the gene expression matrices into a single Seurat object. Cells were automatically annotated using the PBMC reference made available by the Azimuth annotation tool so that non-T cells could be excluded from signature analysis^128^. Cells receiving CD4 T, CD8 T, as well as NK labels were kept, because T cells occasionally receive NK cell labels with automated cell annotation methods. Markers distinguishing MED12-knockout cells from AAVS1-knockout cells were identified using the logistic regression test from the FindMarkers function (min.pct=0.1, logfc.threshold=0.25, only.pos=TRUE). The top 50 markers were selected for module scoring both the Perturb-seq and CRISPR-All-seq objects. In summary, the LTBR signature is computed using the CRISPR-All-seq cells for both the CRISPR-All-seq and OverCITE-seq datasets and the MED12 signature is computed using the Perturb-seq object for both the CRISPR-All-seq and Perturb-seq datasets.

### CRISPR-All-seq Knock-in Sequence Quantification

The codon optimized knock-in sequences are sufficiently distinct from endogenous sequences such that knock-in sequences are not readily aligned to the human genome by the Cellranger pipeline. We initially found that knock-in sequences, such as tNGFR and CAR, are present in gene expression fastqs by minimal ‘grep’ searches and performed further quantification of all knock-in sequences using the kallisto bustools pipeline^129,130^. We first built a ‘reference’ of knock-in sequences which includes the coding sequence of all CAR signaling domains, gene overexpression sequences, and synthetic gene sequence in the pool, as well as the tNGFR and CAR sequences present 5’ of each modification on the same transcripts. Gene knockdown and gene knockout sequences were also included in the reference, however reads failed to align to likely due to the short length of the gRNA and shRNA sequences. The reference was indexed with kallisto. Each lane of the 10x Genomics chip was then separately processed with kallisto bus (specifying 10xv3 tech), bustools correct, bustools sort, and bustools count. Default parameters were used at each step with bustools version 0.41.0 and kallisto version 0.46.2. The resulting counts matrix was loaded into the Seurat object using the AddMetaData function so concordance with barcode-derived assignments could be determined.

### CRISPR-All-seq Perturbation Assignment Summary Statistics

Summary statistics for CRISPR-All-seq perturbation assignments were computed by subsetting to the fraction of cells where a barcode based assignment and a construct sequence-based assignment was determined. Construct sequence assignments were determined by assigning cells to whichever unique construct is expressed and excluding all cells where more than one unique construct is detected. This differs from the barcode based assignments using deMULTIplex2, because the Construct sequence matrix is far sparser resulting in poorly fitted assignment models. Thus, a simplistic assignment approach was used. The construct sequence assignments are treated as the ground truth assignments because they represent direct capture of the sequence, and barcodes are susceptible to swapping events during amplification and pooled cloning.

### CRISPR-All-seq RNA fingerprinting

RNA fingerprinting is performed to identify similar or shared perturbation signatures spanning perturbation and cell type^131^. The pre-computed Genome-Wide Perturb-Seq dataset^132^ is downloaded from the RNAFingerprinting and RNAFingerprintingData R packages (versions 0.0.0.900 and 0.1.0) and used as a reference dictionary. Default parameters are used for all steps following the reference example on GitHub: https://github.com/satijalab/rna-fingerprinting.

### Bulk RNA Sequencing and Analysis

Bulk RNA sequencing libraries were sequenced on a NovaSeq X Plus in a 150 x 150 nucleotide paired-end format. Analysis was performed using the Nextflow (version 24.10.5) core ‘rnaseq’ pipeline^133^ (https://zenodo.org/records/17153746). Reads were pseudoaligned to the GRCh38 reference using salmon^134^. Differential expression analysis was performed using DESeq2^135^. DEGs for each construct were computed by contrasting with the control construct (28ζ-GFP-NTgRNA-NTshRNA). Following log2 fold-change shrinkage using the lfcShrink (type = “normal”) function, DEG lists were filtered to genes with greater than a 0.25 log2 fold-change and an adjusted p-value less than 0.01. DEG lists for constructs sharing one or more perturbations were intersected in an effort to quantify combinatorial effects of perturbations. GO terms were computed using the clusterProfiler R package version 4.12.6 using the enrichGO function and a p-value cutoff of 0.05^136^.

## RESOURCE AVAILABILITY

### Lead Contact

Further information and requests for constructs, data availability or resources should be directed to the lead contact, Theodore Roth.

### Materials Availability

All DNA sequences used in this study are listed in **Tables S1-3, S9-15,** and **S18-19**. All plasmid libraries and key individual plasmids will be made available on Addgene upon publication. Please contact for annotated sequence maps or any other reagent requests.

### Data Availability

All long-read sequencing, scRNA-seq and bulk RNA-seq data will be deposited to NIH GEO upon publication. Amplicon sequencing counts, bulk RNA-seq counts, the CRISPR-All-seq Seurat object, and long read sequencing barcode and construct assignments are each publicly available on Zenodo (https://doi.org/10.5281/zenodo.17594332). Long-read sequencing data is available upon request.

### Code Availability

Code to reproduce analyses is publicly available on GitHub (https://github.com/AustinHartman/crispr-all) using the data available from Zenodo as input.

## ACKNOWLEDGEMENTS

We thank the members of the Satpathy and Roth Labs for stimulating discussions. The Roth Lab has received sponsored research support from an NIH Director’s New Innovator Award (DP2 CA311217), the NCI (K08 CA286740), a Burroughs Wellcome Career Award for Medical Scientists, a Senior Fellow award from the Parker Institute of Cancer Immunotherapy, an Innovation Investigator Award from the Arc Institute, a The Cancer League Research Grant Program Award, a Gates Foundation Innovation Pilot Award, and received support from the Weill Foundation and Northpond Ventures. T.L.R. is a Member of the Parker Institute of Cancer Immunotherapy, an Affiliate Investigator of the Arc Institute, and an Investigator at the Weill Cancer Hub West. A.T.S. was supported by a Lloyd J. Old STAR Award from the Cancer Research Institute, the Parker Institute for Cancer Immunotherapy, a Mark Foundation Emerging Leader Award, Yosemite, and Northpond Ventures. This work was supported by the Department of Pathology at Stanford University. We thank all anonymous blood donors for supporting this work.

## AUTHOR CONTRIBUTIONS

A.H., A.T.S., and T.L.R. conceptualized the study. A.H., A.T.S., and T.L.R wrote and edited the manuscript with input from all authors. A.H., O.T., C.K., N.E.T., J.L., L.W., M.M., A.M., N.J., L.M., S.M., L.C.C., F.H., A.E., A.C., L.M., T.R., A.H., and T.L.R. performed experiments. A.H. set up computational pipelines and analyzed the data. A.T.S. and T.L.R. guided experiments and data analysis.

## SUPPLEMENTAL INFORMATION

Table S1. Intron Sequences.

Table S2. CRISPR-All format annotated sequences for each perturbation function.

Table S3. x6 and x10 CRISPR-All format combinatorial perturbation DNA sequences.

Table S4. CRISPR-All format human gene knockout library.

Table S5. CRISPR-All format human gene knockdown library.

Table S6. Engineered T Cell meta-analysis full reference list.

Table S7. Engineered T Cell meta-analysis pooled screens summary.

Table S8. CRISPR-All Cell Therapy Universal Screening (CACTUS) meta-library all targets list.

Table S9. FDA approved CARs CACTUS sub-library sequences.

Table S10. CAR binder Domain CACTUS sub-library sequences.

Table S11. CAR signaling Domain CACTUS sub-library sequences.

Table S12. Natural Gene overexpression CACTUS sub-library sequences.

Table S13. Synthetic Gene CACTUS sub-library sequences.

Table S14. Gene Knockout CACTUS sub-library sequences.

Table S15. Gene Knockdown CACTUS sub-library sequences.

Table S16. CACTUS meta-library barcode table.

Table S17. CACTUS screens barcodes counts matrix.

Table S18. CRISPR-All-seq single cell sub-library sequences and barcodes.

Table S19. CRISPR-All combinatorial perturbation library sequences and barcodes.

Table S20. CRISPR-All combinatorial perturbation library screens counts matrices.

## REFERENCES

1. Naldini, L. et al. In vivo gene delivery and stable transduction of nondividing cells by a lentiviral vector. Science 272, 263–267 (1996).

2. Sadelain, M., Wang, C. H., Antoniou, M., Grosveld, F. & Mulligan, R. C. Generation of a high-titer retroviral vector capable of expressing high levels of the human beta-globin gene. Proc. Natl. Acad. Sci. U. S. A. 92, 6728–6732 (1995).

3. Blaese, R. M. et al. T lymphocyte-directed gene therapy for ADA-SCID: initial trial results after 4 years. Science 270, 475–480 (1995).

4. Musunuru, K. et al. Patient-Specific In Vivo Gene Editing to Treat a Rare Genetic Disease. N. Engl. J. Med. 392, 2235–2243 (2025).

5. Tremblay, J. P., Annoni, A. & Suzuki, M. Three Decades of Clinical Gene Therapy: From Experimental Technologies to Viable Treatments. Mol. Ther. 29, 411–412 (2021).

6. Roth, T. L. & Marson, A. Genetic Disease and Therapy. Annu. Rev. Pathol. 16, 145–166 (2021).

7. Stegmeier, F., Hu, G., Rickles, R. J., Hannon, G. J. & Elledge, S. J. A lentiviral microRNA-based system for single-copy polymerase II-regulated RNA interference in mammalian cells. Proc. Natl. Acad. Sci. 102, 13212–13217 (2005).

8. Cong, L. et al. Multiplex genome engineering using CRISPR/Cas systems. Science 339, 819–823 (2013).

9. Jinek, M. et al. RNA-programmed genome editing in human cells. eLife 2, e00471 (2013).

10. Hsu, P. D. et al. DNA targeting specificity of RNA-guided Cas9 nucleases. Nat. Biotechnol. 31, 827–832 (2013).

11. Xue, C. & Greene, E. C. DNA repair pathway choices in CRISPR-Cas9 mediated genome editing. Trends Genet. TIG 37, 639–656 (2021).

12. Schwank, G. et al. Functional Repair of CFTR by CRISPR/Cas9 in Intestinal Stem Cell Organoids of Cystic Fibrosis Patients. Cell Stem Cell 13, 653–658 (2013).

13. Schumann, K. et al. Generation of knock-in primary human T cells using Cas9 ribonucleoproteins. Proc. Natl. Acad. Sci. 112, 10437–10442 (2015).

14. Roth, T. L. et al. Reprogramming human T cell function and specificity with non-viral genome targeting. Nature 559, 405–409 (2018).

15. Vaidyanathan, S. et al. Targeted replacement of full-length CFTR in human airway stem cells by CRISPR-Cas9 for pan-mutation correction in the endogenous locus. Mol. Ther. 30, 223–237 (2022).

16. Shy, B. R. et al. High-yield genome engineering in primary cells using a hybrid ssDNA repair template and small-molecule cocktails. Nat. Biotechnol. 41, 521–531 (2023).

17. Eyquem, J. et al. Targeting a CAR to the TRAC locus with CRISPR/Cas9 enhances tumour rejection. Nature 543, 113–117 (2017).

18. Chang, C. R. et al. SEED-Selection enables high-efficiency enrichment of primary T cells edited at multiple loci. Nat. Biotechnol. 1–11 (2025) doi:10.1038/s41587-024-02531-6.

19. Roth, T. L. et al. Non-viral intron knock-ins for targeted gene integration into human T cells and for T-cell selection. *Nat*. Biomed. Eng. 9, 1309–1319 (2025).

20. Howden, S. E., Thomson, J. A. & Little, M. H. Simultaneous reprogramming and gene editing of human fibroblasts. Nat. Protoc. 13, 875–898 (2018).

21. Mandal, P. K. et al. Efficient ablation of genes in human hematopoietic stem and effector cells using CRISPR/Cas9. Cell Stem Cell 15, 643–652 (2014).

22. Dever, D. P. et al. CRISPR/Cas9 β-globin gene targeting in human haematopoietic stem cells. Nature 539, 384–389 (2016).

23. Shalem, O. et al. Genome-scale CRISPR-Cas9 knockout screening in human cells. Science 343, 84–87 (2014).

24. Wang, T., Wei, J. J., Sabatini, D. M. & Lander, E. S. Genetic Screens in Human Cells Using the CRISPR-Cas9 System. Science 343, 80–84 (2014).

25. Shifrut, E. et al. Genome-wide CRISPR Screens in Primary Human T Cells Reveal Key Regulators of Immune Function. Cell 175, 1958–1971.e15 (2018).

26. Berns, K. et al. A large-scale RNAi screen in human cells identifies new components of the p53 pathway. Nature 428, 431–437 (2004).

27. Root, D. E., Hacohen, N., Hahn, W. C., Lander, E. S. & Sabatini, D. M. Genome-scale loss-of-function screening with a lentiviral RNAi library. Nat. Methods 3, 715–719 (2006).

28. Silva, J. M. et al. Profiling Essential Genes in Human Mammary Cells by Multiplex RNAi Screening. Science 319, 617–620 (2008).

29. Wong, A. S. L. et al. Multiplexed barcoded CRISPR-Cas9 screening enabled by CombiGEM. Proc. Natl. Acad. Sci. 113, 2544–2549 (2016).

30. Zhou, P. et al. A Three-Way Combinatorial CRISPR Screen for Analyzing Interactions among Druggable Targets. Cell Rep. 32, 108020 (2020).

31. Zhou, P., Wan, Y. K., Chan, B. K. C., Choi, G. C. G. & Wong, A. S. L. Extensible combinatorial CRISPR screening in mammalian cells. STAR Protoc. 2, 100255 (2021).

32. Wong, A. S. L., Choi, G. C. G., Cheng, A. A., Purcell, O. & Lu, T. K. Massively parallel high-order combinatorial genetics in human cells. Nat. Biotechnol. 33, 952–961 (2015).

33. Qi, L. S. et al. Repurposing CRISPR as an RNA-guided platform for sequence-specific control of gene expression. Cell 152, 1173–1183 (2013).

34. Gilbert, L. A. et al. Genome-Scale CRISPR-Mediated Control of Gene Repression and Activation. Cell 159, 647–661 (2014).

35. Schmidt, R. et al. CRISPR activation and interference screens decode stimulation responses in primary human T cells. Science 375, eabj4008 (2022).

36. Konermann, S. et al. Transcriptome Engineering with RNA-Targeting Type VI-D CRISPR Effectors. Cell 173, 665–676.e14 (2018).

37. Wessels, H.-H. et al. Massively parallel Cas13 screens reveal principles for guide RNA design. Nat. Biotechnol. 38, 722–727 (2020).

38. Tieu, V. et al. A versatile CRISPR-Cas13d platform for multiplexed transcriptomic regulation and metabolic engineering in primary human T cells. Cell 187, 1278–1295.e20 (2024).

39. Rual, J.-F. et al. Towards a proteome-scale map of the human protein–protein interaction network. Nature 437, 1173–1178 (2005).

40. Yang, X. et al. A public genome-scale lentiviral expression library of human ORFs. Nat. Methods 8, 659–661 (2011).

41. Arnoldo, A. et al. A genome scale overexpression screen to reveal drug activity in human cells. Genome Med. 6, 32 (2014).

42. Legut, M. et al. A genome-scale screen for synthetic drivers of T cell proliferation. Nature 603, 728–735 (2022).

43. Roth, T. L. et al. Pooled Knockin Targeting for Genome Engineering of Cellular Immunotherapies. Cell 181, 728–744.e21 (2020).

44. Blaeschke, F. et al. Modular pooled discovery of synthetic knockin sequences to program durable cell therapies. Cell 186, 4216–4234.e33 (2023).

45. Gordon, K. S. et al. Screening for CD19-specific chimaeric antigen receptors with enhanced signalling via a barcoded library of intracellular domains. *Nat*. Biomed. Eng. 6, 855–866 (2022).

46. Daniels, K. G. et al. Decoding CAR T cell phenotype using combinatorial signaling motif libraries and machine learning. Science 378, 1194–1200 (2022).

47. Goodman, D. B. et al. Pooled screening of CAR T cells identifies diverse immune signaling domains for next-generation immunotherapies. Sci. Transl. Med. 14, eabm1463 (2022).

48. Castellanos-Rueda, R. et al. speedingCARs: accelerating the engineering of CAR T cells by signaling domain shuffling and single-cell sequencing. Nat. Commun. 13, 6555 (2022).

49. Wu, M.-R. et al. A high-throughput screening and computation platform for identifying synthetic promoters with enhanced cell-state specificity (SPECS). Nat. Commun. 10, 2880 (2019).

50. Choi, G. C. G. et al. Combinatorial mutagenesis en masse optimizes the genome editing activities of SpCas9. Nat. Methods 16, 722–730 (2019).

51. McGee, A. V. et al. Modular vector assembly enables rapid assessment of emerging CRISPR technologies. Cell Genomics 4, (2024).

52. Dixit, A. et al. Perturb-seq: Dissecting molecular circuits with scalable single cell RNA profiling of pooled genetic screens. Cell 167, 1853–1866.e17 (2016).

53. Adamson, B. et al. A Multiplexed Single-Cell CRISPR Screening Platform Enables Systematic Dissection of the Unfolded Protein Response. Cell 167, 1867–1882.e21 (2016).

54. Datlinger, P. et al. Pooled CRISPR screening with single-cell transcriptome read-out. Nat. Methods 14, 297–301 (2017).

55. Wong, A. S. L., Choi, G. C. G. & Lu, T. K. Deciphering Combinatorial Genetics. Annu. Rev. Genet. 50, 515–538 (2016).

56. Schraivogel, D., Steinmetz, L. M. & Parts, L. Pooled Genome-Scale CRISPR Screens in Single Cells. Annu. Rev. Genet. 57, 223–244 (2023).

57. Park, B.-S., Lee, M., Kim, J. & Kim, T. Perturbomics: CRISPR–Cas screening-based functional genomics approach for drug target discovery. Exp. Mol. Med. 57, 1443–1454 (2025).

58. Anglada-Girotto, M., Miravet-Verde, S. & Serrano, L. Using single-cell perturbation screens to decode the regulatory architecture of splicing factor programs. Nucleic Acids Res. 53, gkaf855 (2025).

59. Menon, A. V., et al. Unraveling the future of genomics: CRISPR, single-cell omics, and the applications in cancer and immunology. Front. Genome Ed. 7, (2025).

60. Liu, X. et al. CRISPR-Cas9-mediated multiplex gene editing in CAR-T cells. Cell Res. 27, 154–157 (2017).

61. Ren, J. et al. A versatile system for rapid multiplex genome-edited CAR T cell generation. Oncotarget 8, 17002–17011 (2017).

62. Stadtmauer, E. A. et al. CRISPR-engineered T cells in patients with refractory cancer. Science 367, eaba7365 (2020).

63. Mikkelsen, N. S. et al. Orthogonal CRISPR systems for targeted integration and multiplex base editing enable nonviral engineering of allogeneic CAR-T cells. Mol. Ther. https://doi.org/10.1016/j.ymthe.2025.08.032 (2025) doi:10.1016/j.ymthe.2025.08.032.

64. Gao, Y. et al. Complex transcriptional modulation with orthogonal and inducible dCas9 regulators. Nat. Methods 13, 1043–1049 (2016).

65. Kampmann, M. CRISPRi and CRISPRa screens in mammalian cells for precision biology and medicine. ACS Chem. Biol. 13, 406–416 (2018).

66. Boettcher, M. et al. Dual gene activation and knockout screen reveals directional dependencies in genetic networks. Nat. Biotechnol. 36, 170–178 (2018).

67. Truong, V. A. et al. CRISPRai for simultaneous gene activation and inhibition to promote stem cell chondrogenesis and calvarial bone regeneration. Nucleic Acids Res. 47, e74 (2019).

68. McCarty, N. S., Graham, A. E., Studená, L. & Ledesma-Amaro, R. Multiplexed CRISPR technologies for gene editing and transcriptional regulation. Nat. Commun. 11, 1281 (2020).

69. Hsiung, C. C.-S. et al. Engineered CRISPR-Cas12a for higher-order combinatorial chromatin perturbations. Nat. Biotechnol. 43, 369–383 (2025).

70. Pacalin, N. M. et al. Bidirectional epigenetic editing reveals hierarchies in gene regulation. Nat. Biotechnol. 43, 355–368 (2025).

71. Jafarzadeh, L., Smaani, A. & Delisle, J.-S. Multiplex engineering and multifunction T cells for precise and effective immunotherapies. Front. Immunol. 16, (2025).

72. Goudy, L. et al. Integrated epigenetic and genetic programming of primary human T cells. Nat. Biotechnol. 1–12 (2025) doi:10.1038/s41587-025-02856-w.

73. Schmidt, R. et al. Base-editing mutagenesis maps alleles to tune human T cell functions. Nature 625, 805–812 (2024).

74. Datlinger, P. et al. Systematic discovery of CRISPR-boosted CAR T cell immunotherapies. Nature 646, 963–972 (2025).

75. Dai, X. et al. One-step generation of modular CAR-T cells with AAV–Cpf1. Nat. Methods 16, 247–254 (2019).

76. Dai, X. et al. Massively parallel knock-in engineering of human T cells. Nat. Biotechnol. 41, 1239–1255 (2023).

77. Rood, J. E., Hupalowska, A. & Regev, A. Toward a foundation model of causal cell and tissue biology with a Perturbation Cell and Tissue Atlas. Cell 187, 4520–4545 (2024).

78. Antonsson, S. E. & Melsted, P. Batch correction methods used in single-cell RNA sequencing analyses are often poorly calibrated. Genome Res. 35, 1832–1841 (2025).

79. June, C. H. & Sadelain, M. Chimeric Antigen Receptor Therapy. N. Engl. J. Med. 379, 64–73 (2018).

80. Sterner, R. C. & Sterner, R. M. CAR-T cell therapy: current limitations and potential strategies. Blood Cancer J. 11, 69 (2021).

81. Labanieh, L. & Mackall, C. L. CAR immune cells: design principles, resistance and the next generation. Nature 614, 635–648 (2023).

82. Rupp, L. J. et al. CRISPR/Cas9-mediated PD-1 disruption enhances anti-tumor efficacy of human chimeric antigen receptor T cells. Sci. Rep. 7, 737 (2017).

83. Lynn, R. C. et al. c-Jun overexpression in CAR T cells induces exhaustion resistance. Nature 576, 293–300 (2019).

84. Xie, S., Cooley, A., Armendariz, D., Zhou, P. & Hon, G. C. Frequent sgRNA-barcode recombination in single-cell perturbation assays. PLoS ONE 13, e0198635 (2018).

85. Sack, L. M., Davoli, T., Xu, Q., Li, M. Z. & Elledge, S. J. Sources of Error in Mammalian Genetic Screens. G3 Bethesda Md 6, 2781–2790 (2016).

86. Hanna, R. E. & Doench, J. G. A case of mistaken identity. Nat. Biotechnol. 36, 802–804 (2018).

87. Hegde, M., Strand, C., Hanna, R. E. & Doench, J. G. Uncoupling of sgRNAs from their associated barcodes during PCR amplification of combinatorial CRISPR screens. PLOS ONE 13, e0197547 (2018).

88. Feldman, D., Singh, A., Garrity, A. J. & Blainey, P. C. Lentiviral co-packaging mitigates the effects of intermolecular recombination and multiple integrations in pooled genetic screens. 262121 Preprint at 10.1101/262121 (2018).

89. Bock, C. et al. High-content CRISPR screening. Nat. Rev. Methods Primer 2, 8 (2022).

90. Ng, A. H. M. et al. A comprehensive library of human transcription factors for cell fate engineering. Nat. Biotechnol. 39, 510–519 (2021).

91. Cheng, A. A., Ding, H. & Lu, T. K. Enhanced killing of antibiotic-resistant bacteria enabled by massively parallel combinatorial genetics. Proc. Natl. Acad. Sci. 111, 12462–12467 (2014).

92. Hernandez Hernandez, D., et al. Improved Combinatorial Assembly and Barcode Sequencing for Gene-Sized DNA Constructs. ACS Synth. Biol. 12, 2778–2782 (2023).

93. Zetsche, B. et al. Cpf1 Is a Single RNA-Guided Endonuclease of a Class 2 CRISPR-Cas System. Cell 163, 759–771 (2015).

94. Fellmann, C. et al. An optimized microRNA backbone for effective single-copy RNAi. Cell Rep. 5, 1704–1713 (2013).

95. Leibowitz, M. L. et al. Chromothripsis as an on-target consequence of CRISPR–Cas9 genome editing. Nat. Genet. 53, 895–905 (2021).

96. Campa, C. C., Weisbach, N. R., Santinha, A. J., Incarnato, D. & Platt, R. J. Multiplexed genome engineering by Cas12a and CRISPR arrays encoded on single transcripts. Nat. Methods 16, 887–893 (2019).

97. McCarty, N. S., Graham, A. E., Studená, L. & Ledesma-Amaro, R. Multiplexed CRISPR technologies for gene editing and transcriptional regulation. Nat. Commun. 11, 1281 (2020).

98. Petiwala, S. et al. Optimization of Genomewide CRISPR Screens Using AsCas12a and Multi-Guide Arrays. CRISPR J. 6, 75–82 (2023).

99. Pelossof, R. et al. Prediction of potent shRNAs with a sequential classification algorithm. Nat. Biotechnol. 35, 350–353 (2017).

100. Labanieh, L. & Mackall, C. L. CAR immune cells: design principles, resistance and the next generation. Nature 614, 635–648 (2023).

101. Tao, R. et al. Revolutionizing cancer treatment: enhancing CAR-T cell therapy with CRISPR/Cas9 gene editing technology. Front. Immunol. 15, 1354825 (2024).

102. Belk, J. A. et al. Genome-wide CRISPR screens of T cell exhaustion identify chromatin remodeling factors that limit T cell persistence. Cancer Cell 40, 768–786.e7 (2022).

103. Dai, X. et al. Massively parallel knock-in engineering of human T cells. Nat. Biotechnol. 41, 1239–1255 (2023).

104. Wu, J. E. et al. In vitro modeling of CD8+ T cell exhaustion enables CRISPR screening to reveal a role for BHLHE40. Sci. Immunol. 8, eade3369 (2023).

105. Rios, X. et al. Refining chimeric antigen receptors via barcoded protein domain combination pooled screening. Mol. Ther. J. Am. Soc. Gene Ther. 31, 3210–3224 (2023).

106. Takacsi-Nagy, O. et al. Evolutionarily guided transcription factor design programs novel T cell states. 2024.11.06.622344 Preprint at 10.1101/2024.11.06.622344 (2024).

107. Dotti, G. et al. Human cytotoxic T lymphocytes with reduced sensitivity to Fas-induced apoptosis. Blood 105, 4677–4684 (2005).

108. Daniel-Meshulam, I., Horovitz-Fried, M. & Cohen, C. J. Enhanced antitumor activity mediated by human 4-1BB-engineered T cells. Int. J. Cancer 133, 2903–2913 (2013).

109. Dong, M. B. et al. Systematic Immunotherapy Target Discovery Using Genome-Scale In Vivo CRISPR Screens in CD8 T Cells. Cell 178, 1189–1204.e23 (2019).

110. Freitas, K. A. et al. Enhanced T cell effector activity by targeting the Mediator kinase module. Science 378, eabn5647 (2022).

111. Arce, M. M. et al. Central control of dynamic gene circuits governs T cell rest and activation. Nature 637, 930–939 (2025).

112. Wong, A. S. L. et al. Multiplexed barcoded CRISPR-Cas9 screening enabled by CombiGEM. Proc. Natl. Acad. Sci. U. S. A. 113, 2544–2549 (2016).

113. Shen, J. P. et al. Combinatorial CRISPR-Cas9 screens for de novo mapping of genetic interactions. Nat. Methods 14, 573–576 (2017).

114. Li, R. et al. Comparative optimization of combinatorial CRISPR screens. Nat. Commun. 13, 2469 (2022).

115. Rood, J. E., Hupalowska, A. & Regev, A. Toward a foundation model of causal cell and tissue biology with a Perturbation Cell and Tissue Atlas. Cell 187, 4520–4545 (2024).

116. Szałata, A. et al. Transformers in single-cell omics: a review and new perspectives. Nat. Methods 21, 1430–1443 (2024).

117. Hao, M. et al. Large-scale foundation model on single-cell transcriptomics. Nat. Methods 21, 1481–1491 (2024).

118. Reese, M. G., Eeckman, F. H., Kulp, D. & Haussler, D. Improved splice site detection in Genie. J. Comput. Biol. J. Comput. Mol. Cell Biol. 4, 311–323 (1997).

119. Li, H. Minimap2: pairwise alignment for nucleotide sequences. Bioinformatics 34, 3094–3100 (2018).

120. Li, H. lh3/bioawk. (2025).

121. Zheng, G. X. Y. et al. Massively parallel digital transcriptional profiling of single cells. Nat. Commun. 8, 14049 (2017).

122. Hao, Y. et al. Dictionary learning for integrative, multimodal and scalable single-cell analysis. Nat. Biotechnol. 42, 293–304 (2024).

123. Zhu, Q., Conrad, D. N. & Gartner, Z. J. deMULTIplex2: robust sample demultiplexing for scRNA-seq. Genome Biol. 25, 37 (2024).

124. Korsunsky, I. et al. Fast, sensitive and accurate integration of single-cell data with Harmony. Nat. Methods 16, 1289–1296 (2019).

125. Korsunsky, I., Nathan, A., Millard, N. & Raychaudhuri, S. Presto scales Wilcoxon and auROC analyses to millions of observations. 653253 Preprint at 10.1101/653253 (2019).

126. Ritchie, M. E. et al. limma powers differential expression analyses for RNA-sequencing and microarray studies. Nucleic Acids Res. 43, e47 (2015).

127. Barrett, T. et al. NCBI GEO: archive for functional genomics data sets—update. Nucleic Acids Res. 41, D991–D995 (2013).

128. Hao, Y. et al. Integrated analysis of multimodal single-cell data. Cell 184, 3573–3587.e29 (2021).

129. Melsted, P. et al. Modular, efficient and constant-memory single-cell RNA-seq preprocessing. Nat. Biotechnol. 39, 813–818 (2021).

130. Bray, N. L., Pimentel, H., Melsted, P. & Pachter, L. Near-optimal probabilistic RNA-seq quantification. Nat. Biotechnol. 34, 525–527 (2016).

131. Grabski, I. N. et al. Mapping transcriptional responses to cellular perturbation dictionaries with RNA fingerprinting. 2025.09.19.676866 Preprint at 10.1101/2025.09.19.676866 (2025).

132. Replogle, J. M. et al. Mapping information-rich genotype-phenotype landscapes with genome-scale Perturb-seq. Cell 185, 2559–2575.e28 (2022).

133. Ewels, P. A. et al. The nf-core framework for community-curated bioinformatics pipelines. Nat. Biotechnol. 38, 276–278 (2020).

134. Patro, R., Duggal, G., Love, M. I., Irizarry, R. A. & Kingsford, C. Salmon provides fast and bias-aware quantification of transcript expression. Nat. Methods 14, 417–419 (2017).

135. Love, M. I., Huber, W. & Anders, S. Moderated estimation of fold change and dispersion for RNA-seq data with DESeq2. Genome Biol. 15, 550 (2014).

136. Yu, G., Wang, L.-G., Han, Y. & He, Q.-Y. clusterProfiler: an R Package for Comparing Biological Themes Among Gene Clusters. OMICS J. Integr. Biol. 16, 284–287 (2012).

